# Reverse transcriptase enzyme and priming strategy affect quantification and diversity of environmental transcripts

**DOI:** 10.1101/2020.03.18.996603

**Authors:** Fabien Cholet, Umer Z. Ijaz, Cindy J. Smith

## Abstract

RT-Q-PCR, and RT-PCR amplicon sequencing, provide a convenient, target-specific, high-sensitivity approach for gene expression studies and are widely used in environmental microbiology. Yet, the effectiveness and reproducibility of the reverse transcription step has not been evaluated. Therefore, we tested a combination of four commercial reverse transcriptases with two priming techniques to faithfully transcribe *16S rRNA* and *amoA* transcripts from marine sediments. Both enzyme and priming strategy greatly affected quantification of the exact same target with differences of up to 600-fold. Furthermore, the choice of RT system significantly changed the communities recovered. For *16S rRNA*, both enzyme and priming had a significant effect with enzyme having a stronger impact than priming. Inversely, for *amoA* only the change in priming strategy resulted in significant differences between the same sample. Specifically, more OTUs and better coverage of *amoA* transcripts diversity were obtained with GS priming indicating this approach was better at recovering the diversity of *amoA* transcripts. Moreover, sequencing of RNA mock communities revealed that, even though transcript α diversities (*i.e.* OTU counts within a sample) can be biased by the RT, the comparison of β diversities (*i.e.* differences in OTU counts between samples) is reliable as those biases are reproducible between environments.

**Originality-Significance Statement:** Is the complementary DNA (cDNA) produced after Reverse Transcription (RT) a faithful representation of the starting RNA? This is a fundamental and important question for transcriptomic-based studies in environmental microbiology that aim to quantify and/or examine the diversity of transcripts via RT approaches. Yet little is known about the reliability and reproducibility of this step. Here, we evaluated the effect of the two main components of the RT reaction – the retro transcriptase enzyme and priming strategy (gene specific vs random priming), on the quantification and diversity of cDNA. We found that both have a significant impact. We further provide evidence to enable informed choices as to the enzyme and priming combinations to improve the performance of RT-PCR approaches. Taken together, this work will improve the reliability and reproducibility of transcript-based studies in environmental microbiology.

## Introduction

Whereas modifications of the genome can reflect adaptations of living organisms over evolutionary time scales, changes in the transcriptome reflect short term responses of cells (López-Maury *et al*., 2008; Browning and Busby, 2016). In environmental microbiology, transcriptomics is essential to understanding which biochemical pathways are triggered by environmental conditions at a given time. RNAseq approaches facilitate primer free metatranscriptomics to reveal global gene expression profiles. It is now a widely used method in environmental microbiology and has allowed scientists to gain formidable insight into the genome-scale mechanisms used by microbes to adapt to changing environmental conditions (Shakya *et al*., 2013; Gutleben *et al*., 2018). However, it generally comes at high cost and requires extensive data analysis. Plus, as an untargeted approach, it may require enrichment of the mRNA (via removal of ribosomal RNA) and will be dependent on sequencing depth to reveal rare transcripts among the diverse array of transcripts expressed in complex environmental samples. In contrast Reverse-Transcriptase-Quantitative PCR (RT-Q-PCR) is directed via primers towards a single target. While this is much lower throughput in terms of a global overview of transcription, this approach facilitates transcript quantification that is specific, with high-sensitivity and low-detection limits over a wide dynamic range (Sanders *et al*., 2014). RT-Q-PCR is high-throughput in terms of sample numbers, cost effective (in comparison to metatranscriptomics) and subsequent data processing is fast without the requirement for high computational power and bioinformatic expertise needed for metatranscriptomics analysis. As a consequence, RT-(Q)-PCR is routinely used in most life science research fields including environmental microbiology to target and quantify specific transcripts.

In environmental microbiology it is an invaluable approach to further link microbial activity, via gene expression, to microbial and ecosystem processes, compared to DNA approaches alone (Smith and Osborn, 2009; Saleh-Lakha *et al*., 2011; Gadkar and Filion, 2013). As a result the approach has been used to quantify transcripts to distinguish different pathways of the nitrogen cycle in sediments (Santoro *et al*., 2010; Smith *et al*., 2007; Zheng *et al*., 2013; Damashek *et al*., 2015; Duff *et al*., 2017; Santos *et al*., 2018; Zhang *et al*., 2018), soil (Leininger *et al*., 2006; Graham *et al*., 2011; Wang *et al*., 2012; Li *et al*., 2017; Pierre *et al*., 2017), water column (Tolar *et al*., 2016; Santoro *et al*., 2010; Kapoor *et al*., 2015; Posman *et al*., 2017; Feng *et al*., 2018; Gonçalves *et al*., 2018; Liu *et al*., 2018; Christiansen *et al*., 2019) and other microbial processes including water treatment (Gadkar and Filion, 2013; Botes *et al*., 2013; Wang *et al*., 2016; Pelissari *et al*., 2017, 2018) and bioremediation (Gadkar and Filion, 2013; Marzorati et *al.,* 2010; Yergeau *et al*., 2009). In addition to this, cDNA from mRNA or rRNA can undergo PCR for amplicon sequencing to reveal actively transcribing organisms within the environment (Cholet *et al.,* 2019; Zhang *et al.,* 2018; Duff *et al.,* 2017). The RT-(Q)-PCR workflow consists of two steps, the RT reaction that converts RNA to cDNA and the subsequent PCR amplification of the cDNA. cDNA can be directly quantified via RT-Q-PCR or undergo end-point PCR for amplicon sequencing of the expressed transcripts. The initial RT reaction requires a reverse transcriptase enzyme, of which there are a number commercially available, and a reverse primer to initiate the RT reaction. There are two main priming strategies, random or gene specific priming. For random priming, short oligonucleotides (e.g. hexamer or decamer) consisting of all possible sequence combinations for that size, are used to randomly initiate the RT across the entire transcriptome. Gene specific, as the name implies, target specific transcripts of interest.

The aim for the RT reaction is that it faithfully produces cDNA that reflects gene expression in the starting RNA sample (Bustin and Nolan, 2017). However, a number of studies in the wider field of molecular biology indicate that the RT reaction has a significant impact on the final results for the same sample. Indeed within clinical studies, the inherent variability of cDNA synthesis has been reported in some cases to be greater than the differences between biological samples (Sanders *et al*., 2014). This level of variability implies that comparison of results between different studies using different approaches is near impossible (Bustin, 2002). Moreover, the sources of RT variability have been attributed to a wide range of factors including: the choice of reverse transcriptase (Ståhlberg *et al*., 2004; Stangegaard *et al*., 2006; Levesque-Sergerie *et al*., 2007; Werbrouck *et al*., 2007; Okello *et al*., 2010; Sieber *et al*., 2010; Miranda and Steward, 2017); priming (Lekanne Deprez *et al*., 2002; Ståhlberg *et al*., 2004; Stangegaard *et al*., 2006; Werbrouck *et al*., 2007; Sieber *et al*., 2010; Miranda and Steward, 2017); background RNA concentration (Bustin & Nolan, 2004; Levesque-Sergerie et al., 2007; Miranda & Steward, 2017); cleaning of the RT reaction (Okello *et al*., 2010); RNaseH treatment (Polumuri et al., 2002); RT reaction composition and conditions (Ståhlberg *et al*., 2004; Werbrouck *et al*., 2007) and dilution of cDNA (Smith et al 2006).

In environmental microbiology applications, while the subsequent analysis of cDNA by Q-PCR (Smith et al 2006; Smith & Osborn, 2009), and/or PCR for amplicon sequencing (Marotz *et al.,* 2019) has been shown to be robust, reliable and reproducible, the effect of the initial RT reaction on quantification and amplicon sequencing of environmental transcripts has yet to be determined. Indeed environmental samples may provide a number of further challenges for efficient and reproducible RT reactions due to the presence of co-extracted inhibitors; variable target expression (high to low) in a background of high non-target template concentration and low RNA quality and integrity (Cholet *et al*., 2019). Moreover, there is the need for the RT reaction to faithfully transcribe the diversity of target of interest. A small number of studies investigating primer-free approaches to characterise *16S rRNA* transcripts revealed better accuracy (Mäki and Tiirola, 2018) and sensitivity (Hoshino and Inagaki, 2013) with primer-free approaches for amplicon sequencing than PCR of the cDNA. Nonetheless, these primer-free approaches still rely on an initial RT reaction, which could impact the outcome of the *16S rRNA* transcript sequencing. Moreover, our own personal observations in the laboratory have indicated that RT enzyme and priming strategy greatly impact the results of environmental transcript studies, often meaning the difference between detection or not of a given transcript that in turn results in different ecological interpretation. Therefore, to improve reproducibility and inform best practice and standardisation of RT-(Q)-PCR approaches in environmental microbiology, we have undertaken a detailed study of the effect of the RT reaction on RNA extracted from environmental samples. We aimed to determine the impact of enzyme and priming strategy on quantification and amplicon sequencing of transcripts (spiked artificial RNA, *16S rRNA* and ammonia monooxygenase (*amoA*)). We therefore examined a combination of four commonly used commercial reverse transcriptases (Superscript III, Superscript IV, Omniscript and Sensiscript; designated SSIII, SSIV, Omni and Sensi, respectively, thereafter) and two priming strategies (random hexamer and gene specific; designated RH and GS, respectively, thereafter). We hypothesized that both quantification and alpha diversity (*i.e.* OTU counts within a sample) of transcripts from the same samples will be affected by RT enzyme and priming strategy.

## Results

### Effect of enzyme and priming on the detection of exogenous spike

First, the impact of enzyme and priming on the quantification of RNA was determined for an exogenous target that was spiked at known concentrations into a background of environmental RNA. Artificial RNA (*sfGFP* RNA) that could be distinguished from environmental RNA background, was produced by *in vitro* transcription of a PCR product amplified from the pTHSSd_8 plasmid (Segall-Shapiro *et al*., 2014). The resulting RNA was mixed with environmental RNA at different concentrations (10^3^, 5 x 10^3^, 2 x 10^6^ and 10^7^ copies/µl) and quantified using RT-Q-PCR (Figure 1A).

**Figure 1.**
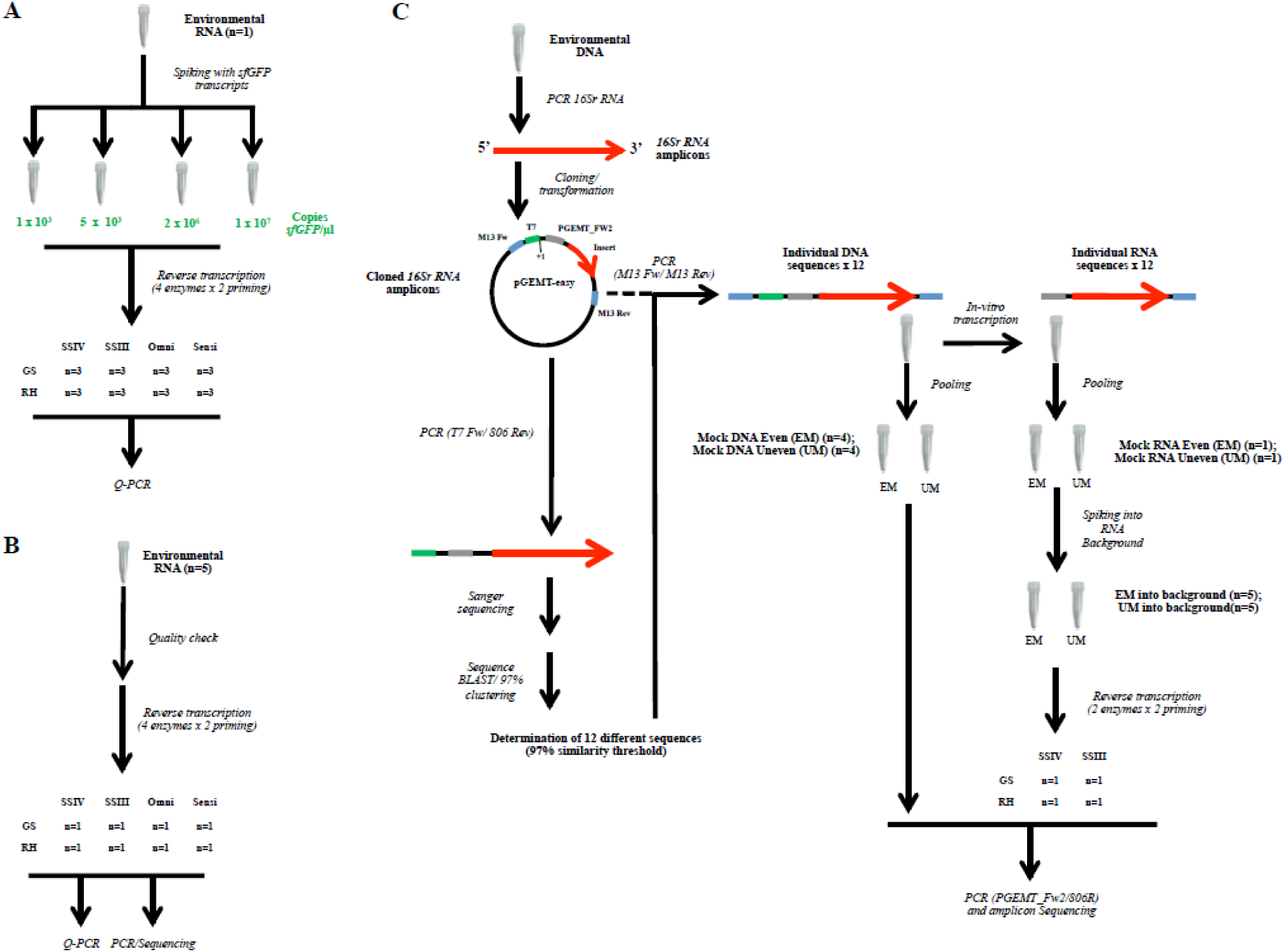
Schematic representation of the experimental workflow followed in this study. The effect of the RT reaction was evaluated on: A) the quantification of an exogenous transcript spiked at known concentrations; B) the quantification of two endogenous transcripts and the subsequent sequencing of these transcripts and C) the sequencing of mock communities composed of 12 transcripts with known sequences for this last experiment, DNA mocks were also included as controls. Replicates are indicated by “n=”.

### sfGFP standard curves

Artificial RNA (*sfGFP* RNA) that could be distinguished from environmental RNA, was produced by *in vitro* transcription of a PCR product amplified from the pTHSSd_8 plasmid. Standard curves were constructed using 10-fold dilutions of *sfGFP* RNA from 10^10^ to 10^1^ copies/µl that underwent individual reverse transcription in duplicate using each of the four RT enzymes with two priming strategies. The Cycle thresholds (Cts) of the same sample derived from different enzyme/priming strategies were obtained by Q-PCR amplification of the resulting cDNA. Amplification of the no template controls and log_10_[*sfGFP*]= 1 and 2 gave no signal. The Limit of Detection (LD) and Quantification (Forootan *et al*., 2017) was log_10_[*sfGFP*]= 3 for all RT systems except for Sensi-RH which had a LD at log_10_[*sfGFG*]= 4. Excluding the Cts obtained for log_10_[*sfGFP*]=10 resulted in an improvement of the regression fit (slopes closer to the expected −3.332 and R squared closer to 1; data not shown). As such standard curves ranged between log_10_[*sfGFP*] = [3;9] (or [4;9] for Sensi-RH). For all enzymes, the use of RH resulted in higher Cts than GS at all *sfGFP* concentrations. SSIV-GS resulted in the best efficiency (99.3%) while the lowest was obtained by Sensi-RH (84.2%) (Table S.1).

Comparison of regressions between RT methods (enzyme/priming) revealed significant difference between slopes (F=3.29; Df=7; *p.*value=0.0036) (GraphPad Prism6, www.graphpad.com). The effect of the RT approach on standard curve construction was further investigated using multilevel linear model analysis. Three different models were tested where 1) intercepts only, 2) slopes only and 3) slopes and intercepts varied between groups (*i.e*. RT method). Models 1 and 2 were then tested against model 3 using an ANOVA. Model 3, allowing for variations in both slopes and intercepts, resulted in a better fit than model 1 (intercept only) or model 2 (slopes) (Table S.2) indicating that the effect of enzyme and priming impacted both the slope (*i.e.* efficiency) and the intercept (*i.e.* signal at No Template Control (NTC)) of the standard curves.

### RT-Q-PCR detection of the RNA sfGFP spike

The exogenous RNA spike, *sfGFP,* was added to environmental RNA at known concentrations and the resulting preparations were reverse-transcribed using the eight different combinations of RT enzymes and priming strategies (Figure 1A). Four different concentrations of spike were added to environmental RNA background: two low and two high, with a five-fold difference in *sfGFP* copy number between the two low and the two high spikes respectively (10^3^ and 5 x10^3^ copies/µl for low spikes; 2 x10^6^ and 10^7^ copies/µl for high spikes). After cDNA synthesis, the spiked target was quantified by Q-PCR and Cycle thresholds (Ct) were converted to copies/µl using standard curves.

Both enzyme and priming had a strong effect on the copy number of exogenous target detected (log10 transformed) at all spike concentrations (Table 1; Figure 2). Overall, SSIII and SSIV enzymes were the closest to the expected value. SSIV was slightly more accurate than SSIII, especially at spike concentrations > 5 x 10^3^. The use of Omni resulted in an underestimation of the spike concentration with factors ≈4 to ≈50 depending on the concentration of the target (higher differences at higher concentrations). Similarly, the use of Sensi also resulted in an underestimation of the exogenous target concentration with factors ≈3 to ≈30 (higher differences at higher concentrations) (Figure 2).

**Figure 2.**
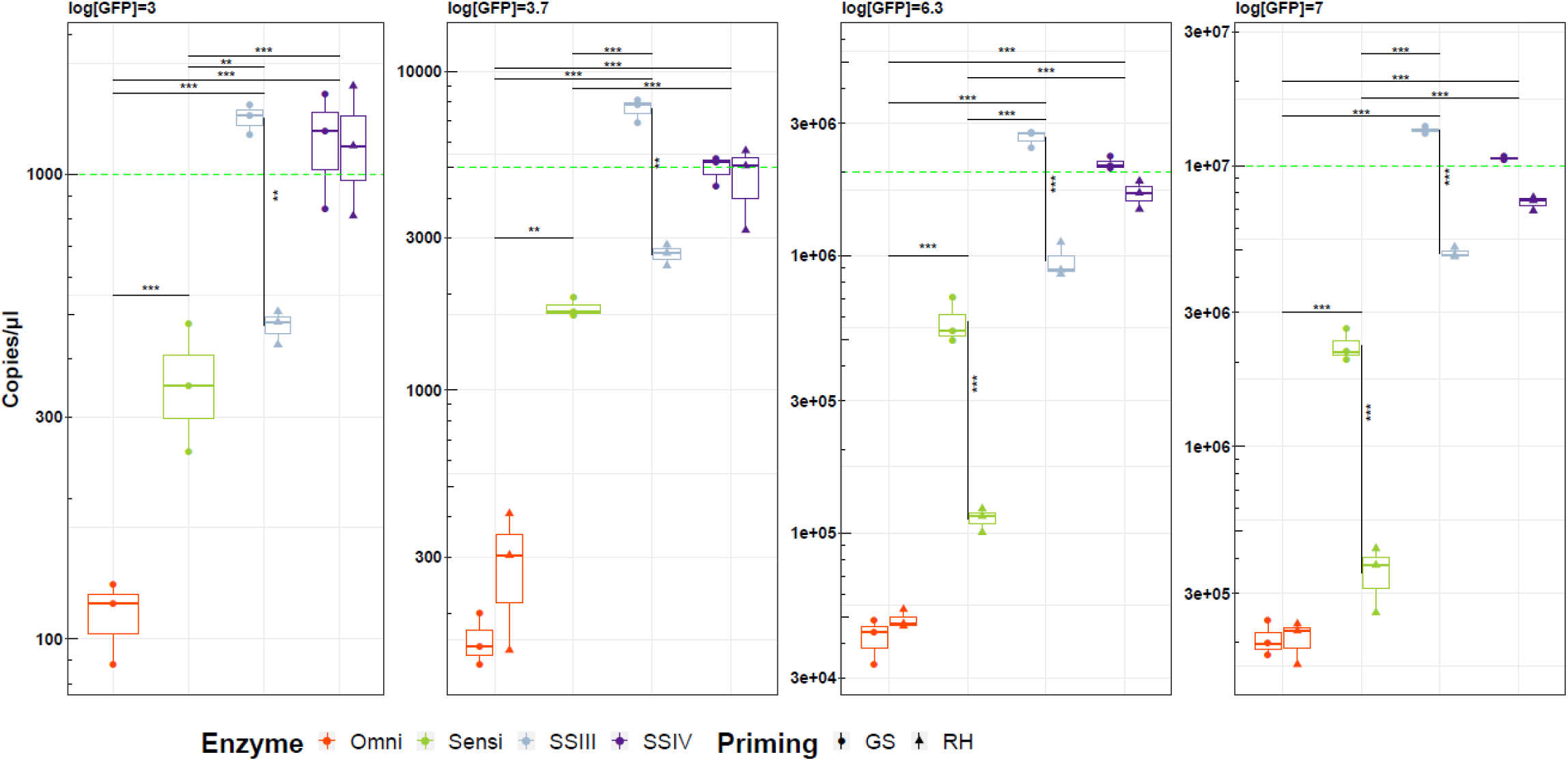
Effect of reverse-transcriptase enzyme and priming on the quantification of the same *sfGFP* spike in environmental RNA background. The concentration of the RNA spike (log[GFP]= 3; 3.7; 6.3and 7) inoculated is indicated at the top of each plot. The results of the two-way ANOVA, showing statistical differences between priming and enzyme for the same template, are presented as vertical and horizontal lines respectively. *: *p*.value <0.05; **: *p*.value <0.01; ***: *p*.value <0.001. The actual concentration of spike for each experiment is indicated by a green dashed line. GS = gene specific; RH = Random Hexamer

**Table 1.** Two-way ANOVA showing the impact of RT system on the quantification of the sfGFP spike.

| Spike concentration (copies/μl) | Enz | Prim | Enz:prim |
| --- | --- | --- | --- |
| 1 x 10<sup>3</sup> | 6.62 x10<sup>-8</sup> | 2.57 x10<sup>-3</sup> | 3.45 x10<sup>-3</sup> |
| 5 x10<sup>3</sup> | 5.92 x10<sup>-12</sup> | 0.08 | 2.94 x10<sup>-4</sup> |
| 2 x 10<sup>6</sup> | <2 x10<sup>-16</sup> | 4.98 x10<sup>-10</sup> | 7.38 x10<sup>-9</sup> |
| 1 x 10<sup>7</sup> | <2 x10<sup>-16</sup> | 1.24 x10<sup>-10</sup> | 1.35 x10<sup>-8</sup> |

*p*.values for the effect of enzyme (Enz), priming (Prim) and the interaction between the two (Enz:Prim) on Ct for the different spike concentrations.

The use of GS priming resulted in more accurate quantification for all enzymes except Omni for which it had no effect. For Sensi, RH priming failed at the low spike concentrations while at the high concentration of spike, RH was significantly lower (6-fold) than GS. For the Superscript enzymes, the use of RH versus GS generally resulted in lower quantification of the same target, except when using SSIV at low concentrations where the priming strategy had no effect. Of the two Superscript enzymes, SSIII with GS always overestimated the concentration of spike whereas RH always underestimated it (≈2 fold or less). Priming had the least effect with SSVI, but more accurate quantification was achieved when using GS priming (Figure 2).

Next, we tested the ability of the RT systems to faithfully report a 5-fold difference in the *sfGFP* spike concentration between the two low and two high concentrations respectively. For this, the differential expression (DE), *i.e.* the ratio of the average transcript number/µl between the two low or the two high spikes respectively, was calculated (Figure 3). The DE does not report how accurately the system quantifies the spike but rather its ability to reflect the 5-fold change. Again, the choice of enzyme and priming had an effect on the observed DE. All systems were better at detecting actual differences (DE closer to 5) at high spike concentrations. The most accurate system, *i.e.* giving DE values closer to the expected 5, at high spike concentration was SSIII-GS (DE = 5.03), followed by SSIV-GS (DE = 4.96) and Omni-GS (DE = 4.91). All enzymes gave less accurate results when used in combination with RH priming at high spike concentration. Still, SSIII and SSIV were the most accurate enzymes, with SSIII better than SSIV. At low spike concentration, SSIII always overestimated the DE whereas SSIV always underestimated it. The use of RH made SSIII slightly more accurate (DE=5.64 with RH versus 5.75 with RH) whereas it made SSIV slightly less accurate (DE=3.94 with RH versus 4.16 with GS). Interestingly, Sensi performed the best at low spike concentration when used with GS priming (DE = 5.03) whereas Omni performed the worst (DE = 1.47). Both Sensi and Omni failed at low spike concentrations when used with RH priming. Therefore, the superscript enzymes preformed best overall, and the DE was improved when SSIII and IV were used in combination with GS priming (Figure 3).

**Figure 3.**
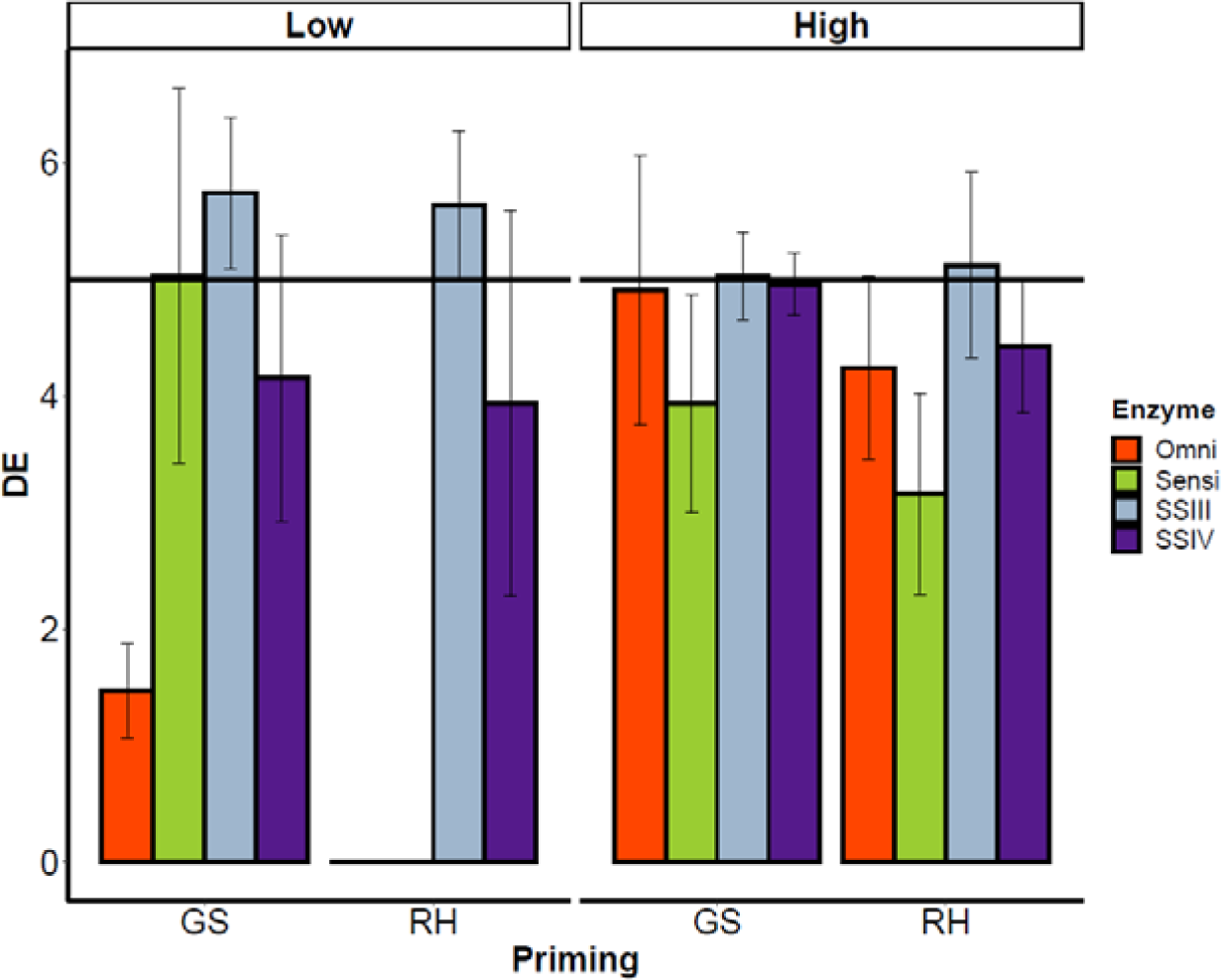
Ability of the RT systems to detect a 5-fold difference of target at low or high concentration in a background of environmental RNA. Differential expression (DE) is the ratio of the average copies/µl between “Low” (1 x 10^3^ and 5 x 10^3^ copies/µl) or “High” (2 x 10^6^ and 1 x 10^7^ copies/µl) *sfGFP* spikes. The expected DE is represented by the horizontal black line at y=5 while the measured DE by each RT system is shown by the bar-plot. GS = gene specific; RH = random hexamer

### Effect of standard curve construction on sfGFP quantification

As there are two approaches to constructing RNA standard curves (Smith *et al.,* 2006), we tested if this had any impact on quantification of the spike and the above results. A standard curve can be made by serial dilution of RNA with individual RTs or via a single RT of RNA followed by serial dilution of cDNA. Standard curves for each enzyme and primer combination were made using these two approaches to quantify the spiked *sfGFP* (Figure 1A). The percentage error was calculated between the observed and expected copies/µl for each *sfGFP* spike generated from each standard curve. The standard curve constructed from the dilution of cDNA generally increased the percentage error, and therefore dilutions of RNA for individual RTs were used for subsequent standard curves (Figure S.1).

### <u>Effect of enzyme and priming on the quantification of endogenous environmental</u> ***<u>transcripts</u>***

#### RNA Quality check

Before proceeding with the quantification of endogenous transcripts, RNA extracted from sediment underwent a quality check (Table 2) (Cholet *et al*., 2019). All samples had good integrity as shown by the RIN (always > 7) and R_amp_ (R_amp_ 380/120 ≈ 0.8 or higher and R_amp_ 380/170 ≈ 0.7 or higher).

**Table 2.** Sediment RNA quality check.

| Sample | Quantity (ng/μl) |  |  | Purity |  |  | Integrity |  |
| --- | --- | --- | --- | --- | --- | --- | --- | --- |
|  | [RNA] | [RNA] | [RNA] | 260/280 | 260/230 | RIN | R<sub>amp</sub> | R<sub>amp</sub> |
|  | Nanodrop | Qubit | Bioanalyser |  |  |  | 380/120 | 380/170 |
| **Env1** | 87.8 | 54.6 | 55.8 | 1.69 | 1.51 | 7.85 | 1.01 | 0.77 |
| **Env2** | 301.2 | 196 | 135 | 1.74 | 2.02 | 7.6 | 0.88 | 0.73 |
| **Env3** | 164 | 117 | 67 | 1.74 | 1.73 | 7.9 | 0.92 | 0.73 |
| **Env4** | 201 | 140 | 91.5 | 1.76 | 1.87 | 7.9 | 0.78 | 0.65 |
| **Env5** | 218.2 | 156 | 91.5 | 1.78 | 1.84 | 7.1 | 0.82 | 0.72 |

Extracted RNA quantity, purity and integrity were determined. RIN = RNA Integrity Number, as determined from Agilent Bioanalyser; R<sub>amp</sub> was calculated as described in Cholet *et al.*, 2019.

#### Quantification of endogenous transcripts

Next, the effect of RT enzyme and priming strategy on the quantification of transcripts from the same sediment sample were tested. For this we targeted *in situ* highly abundant *16S rRNA* and less abundant mRNA from the bacterial ammonia monooxygenase subunit A, *amoA,* for quantification from cDNA generated using the different combinations of reverse transcriptases and priming (Figure 1B). Results were converted into copies/µl using paired standard curves, normalized per µg of extracted RNA and log10 transformed (for parametric 2-ways ANOVA tests). The results clearly showed the effect of the RT system was target dependent: for *16S rRNA* both enzyme and priming significantly affected the results whereas for *amoA* only the effect of enzyme was significant (Figure 4; Table 3).

**Figure 4.**
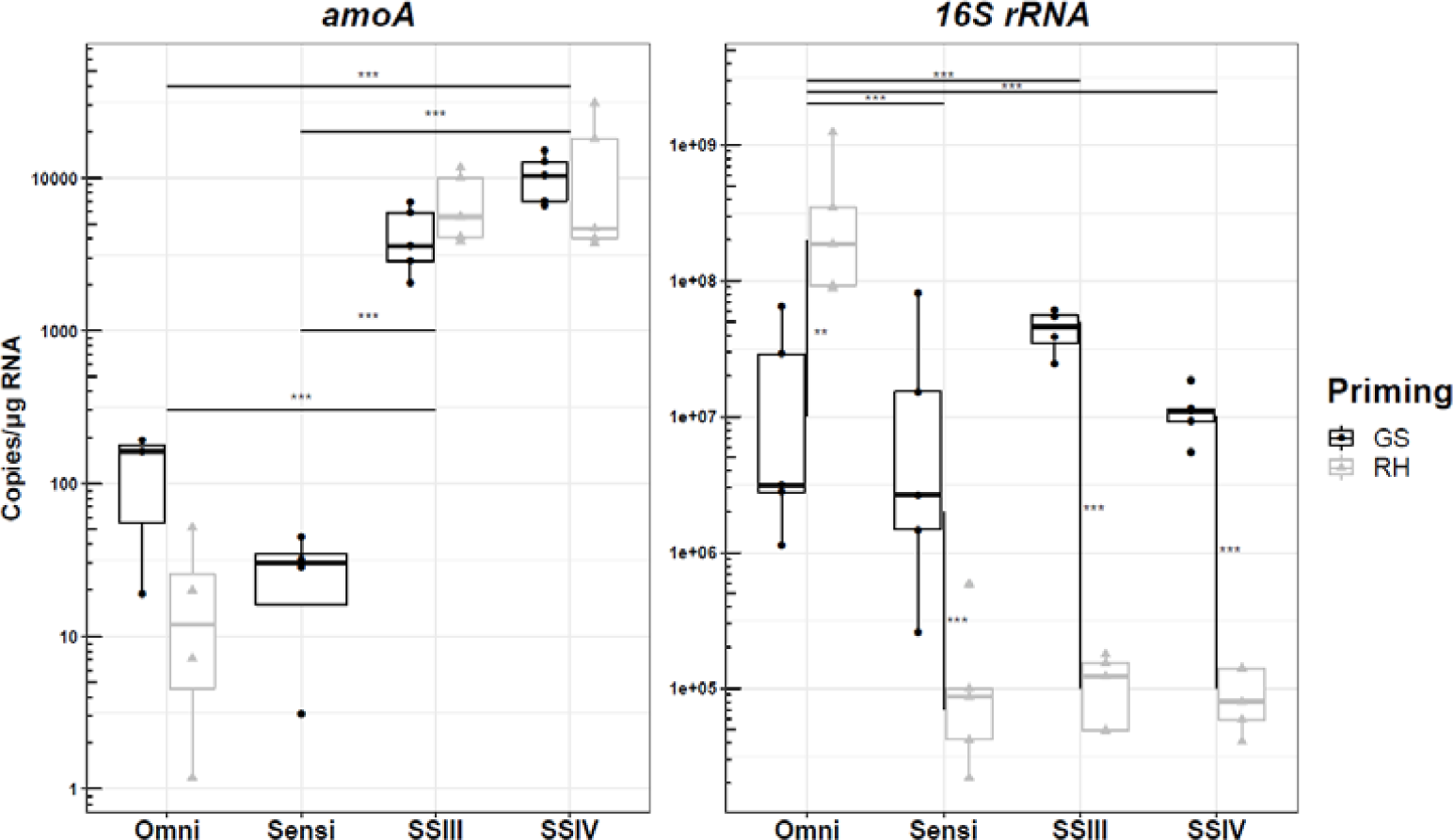
Impact of RT system quantification of two endogenous transcripts from environmental samples. The effect of the RT system on quantification of Bacterial *amoA* (left) and *16S rRNA* (right) transcripts. The results of the two-way ANOVA, showing statistical differences between priming and enzyme for the same template, are presented as vertical and horizontal lines respectively. *: *p*.value<0.05; **: *p*.value<0.01; ***: *p*.value<0.001.

**Table 3.**
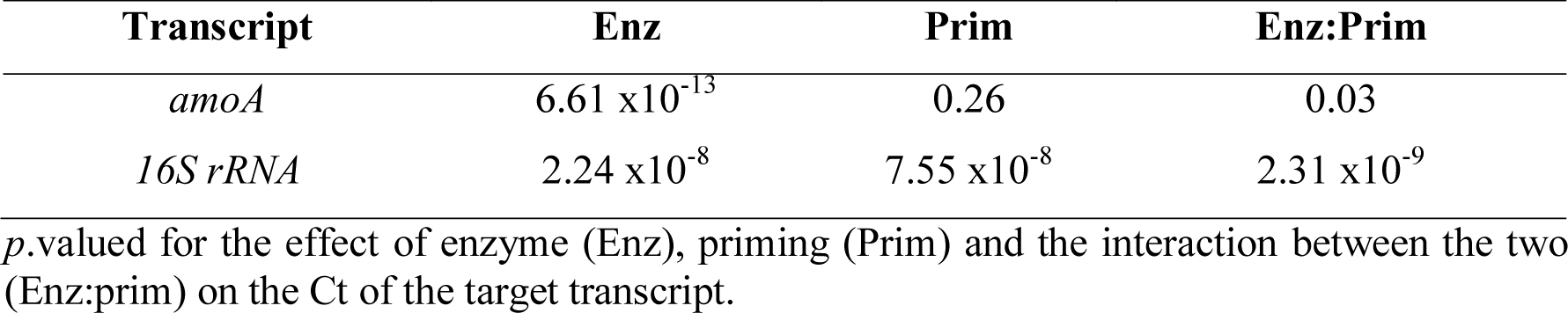
Two-ways ANOVA showing the impact of RT system on the quantification of endogenous transcripts.

| Transcript | Enz | Prim | Enz:Prim |
| --- | --- | --- | --- |
| *amoA* | 6.61 x10<sup>-13</sup> | 0.26 | 0.03 |
| *16S rRNA* | 2.24 x10<sup>-8</sup> | 7.55 x10<sup>-8</sup> | 2.31 x10<sup>-9</sup> |

*p*.valued for the effect of enzyme (Enz), priming (Prim) and the interaction between the two (Enz:prim) on the Ct of the target transcript.

For *amoA*, the choice of RT system resulted in differences of up to 600-fold in the detected copies/µg RNA in the samples tested (Omni-RH versus SSIV-RH) and, in the most extreme case, the difference between detection of the target or not (Sensi). For this assay, only the choice of enzyme significantly affected the results whereas priming did not. A clear difference between Omni/Sensi and the Superscript enzymes (SSIII and SSIV) was observed with, on average, 150 times more *amoA* transcripts per µg RNA with the Superscript enzymes. For Sensi and Omni, the choice of GS priming resulted in better results, especially for Sensi which failed at producing reliable results with RH. For Omni, the use of GS priming resulted in 6 times more copies of *amoA* transcripts compared to RH, although this difference was not statistically significant. SSIII and SSIV performed relatively similarly, with no statistical differences between the two, although the use of SSIV resulted in higher numbers of *amoA* transcripts detected (≈+2.4 fold with GS and ≈+1.7 fold for RH; *p*.value=0.512).

Interestingly, although the use of RH priming resulted in higher Cts on Q-PCR (*i.e.* lower quantification), conversion to copies/µg RNA via the standard curve resulted in a higher quantification than that achieved with GS priming (≈+1.2fold for SSIV (*p*.value=0.99) and ≈+1.7fold for SSIII (*p*.value=0.99)). In summary, SSIV was the best choice for the detection of *amoA* transcripts as it resulted the highest numbers of transcripts detected and produced consistent results between GS and RH. When used in combination with GS as opposed to RH the results were more precise (*i.e.* lower standard deviation) (Figure 4).

For *16S rRNA*, both enzyme and priming had a strong effect on quantification (Figure 4; Table 3). The choice of RT system resulted in differences of ≈ 2300 fold between highest (Omni-RH) and lowest quantification (SSIV-RH > SSIII-RH > Sensi-RH). Omni actually behaved differently from the other enzymes as it was the only one for which the use of RH resulted in higher detected copies/µg RNA compared to GS and indeed, statistical differences were found only between Omni and the other three enzymes. For SSIV, SSIII and Sensi, the use of RH always resulted in lower detected copies *16S rRNA*/µg RNA (≈ −120fold for SSIV and Sensi; ≈ −400fold for SSIII). Results between enzymes were more consistent when used with GS priming, with an average difference in detected copies/µg RNA between enzymes of 2.18-fold (max: 4.01-fold between SSIII and SSIV; min: 1-fold between Sensi and Omni). With this priming, SSIII resulted in the highest number of copies of *16S rRNA*/µg RNA (+4.01fold versus SSIV and +2.22fold versus Omni and Sensi). It is worth mentioning that, even though the use of Omni-GS and Sensi-GS resulted in more copies *16S rRNA*/µg RNA on average compared to SSIV-GS (≈ (*i.e.* lower standard deviation), as was the SSIII-GS combination (Figure 4).

### Effect of enzyme and priming on cDNA amplicon sequencing data

While the quantitative work clearly shows dramatic and significant differences when using different RT enzyme and priming strategies for quantification of the same template, it does not inform if these impact upon community transcript diversity. To examine this, RNA and DNA mock communities of known composition were examined in addition to endogenous *16S rRNA* and *amoA* transcripts from marine sediments.

### Effect on mock community composition

As the actual composition of the transcriptome of the environmental samples is unknown, it is virtually impossible to determine which RT system most closely represents the starting RNA. We thus tested the effect of enzyme and priming on known RNA mock communities. Two mocks community (one even, with all 12 sequences at the same relative proportion, designated EM and one uneven, with the 12 sequences at different relative proportions, designated UM, Table S.3), each composed of twelve different *16S rRNA* transcripts were constructed as detailed in Figure 1C. To further tease apart the effects of the PCR from the RT, both DNA and RNA mock communities were constructed. Of the twelve mock community sequences, one (S9) was over-represented in the DNA mock community but under-represented in the RNA mock community sequencing data. In contrast sequence S10 showed the opposite trend (over-represented in the RNA mock community but under-represented in the DNA mock community). As a result, sequences (S9 & S10) were removed from further analysis.

*DNA mock*. In the EM community, there were variations from the expected proportions (10%). Some members of the community: S1, S2, S4, S8 and S12 were underrepresented whereas S3, S5, S6, S7, S11 were overrepresented. The most underrepresented member, S4, represented only 4% of the total community whereas the most overrepresented, S7, represented 14%. Yet, even though the observed proportions deviated from the expected 10%, they were within the same order of magnitude (Figure 5A).

**Figure 5.**
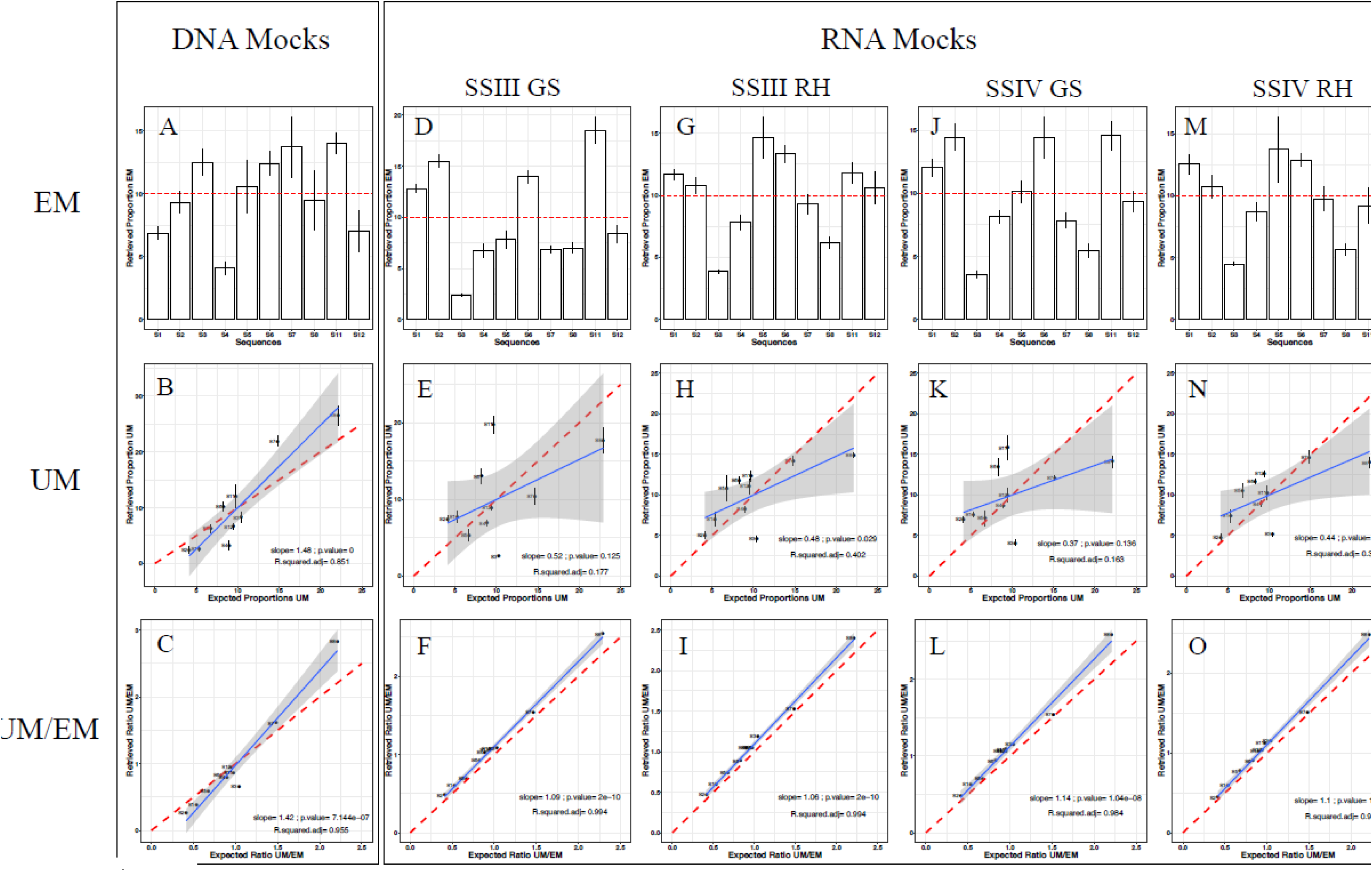
Impact of RT system on the reproducibility of mock community composition. Row EM: observed proportions of each sequence within the Even Mock (EM) communities. The dashed red line indicates the expected proportion. Row UM: regression of observed versus expected proportions of each sequence within the Uneven Mock (UM) communities. Row UM/EM: the observed proportion of each sequence in the UM has been divided by its observed proportion in the EM and plotted against its actual ratio. Row UM and UM/EM the expected regression (y=x) is represented by a dashed red line. The actual regression is represented by a solid blue line with the 95% confidence interval (grey area). Individual regression statistics are reported on individual plot.

For the UM mock communities, the observed proportion of each member was plotted against the expected proportion (Figure 5B). A regression line with equation y=x is expected if each sequence is faithfully represented. The UM community results were consistent with those of the EM, with S6, S7, S11 overrepresented and S1, S2, S4 and S12 underrepresented. S5 was at the correct proportion in both EM and UM.

For most sequences, the errors of representations were consistent between EM and UM (*i.e.* a sequence overrepresented in the EM was generally also over-represented in the UM and vice versa) indicating a sequence specific bias of the PCR/sequencing workflow. And indeed, when the proportions of the UM were corrected by those of the EM, the fit of the regression improved (Figure 5C).

*RNA mock*. As seen for the DNA mock community, the proportions observed in the RNA EM differed from the expected 10% (Figure 5D, G, J, M). When analysing the sequences abundances in the EM using PERMANOVA test (Bray-Curtis distances), it was found that the priming strategy used had a significant effect on the proportions retrieved (*p.*value = 0.001), with RH being more accurate than GS (proportions closer to the expected 10%). On the other hand, neither enzyme nor the interaction between enzyme and priming had a significant effect (*p.*value=0.208 and *p.*value=0.194 respectively). The data set containing the highest errors was SSIII GS, followed by SSIV GS, SSIII RH and the lowest errors were found for SSIV RH (similar amount of error than for the DNA mock) (Figure 5 and Figure S.2). For individual sequences, there was a sequence-specific bias: some sequences (S1, S2 and S6) were always over-represented in the data sets but this was different depending on the priming used (proportion S1 < proportion S2 with GS and inversely with RH). Other members of the mock communities (S3, S4, S7 and S8) had proportions lower than the expected 10%; Though S7 was very close to the expected 10% with SSIV RH. Finally, the results from the other members of the mock community were dependent on the enzyme and priming strategy used. For example, S5 and S12 were always over-represented in the RH-prepared libraries but not in the GS ones. Inversely, S11 was over-represented in the GS libraries (especially with SSIII) and its proportions decreased in the RH ones.

As for the EM, the proportions retrieved for the UM mocks differed from the expected proportions (y=x) (Figure 5E, H, K, N). As for the EM, RH seemed to perform better than GS with better fits for the regressions (for SSIII: R-squared GS = 0.177 versus 0.402 for RH; for SSIV: R-squared GS = 0.163 versus 0.373 for RH). The use of RH priming also resulted in slopes closer to the expected value of 1 compared to GS indicating a better conservation of the relative proportions using this priming strategy.

However, as observed for the DNA mock, the errors were consistent between EM and UM: A sequence over-represented in the EM would also be over-represented in the UM and inversely. As a consequence, when UM reads were corrected by EM reads (Figure 5 F, I, L and O), the calculated slopes were very close to the expected value (y=x) and the R-squared values also improved (close to 1) indicating a better fit of the regression. This observation indicates that the same sequences were misrepresented in both EM and UM communities.

### Effect of RT enzyme and priming strategy on endogenous community composition

*16S rRNA* and (Bacterial) *amoA* PCR amplicons were generated from cDNA prepared using the different combinations of enzymes and priming (Figure 1B). For *amoA*, only SSIII and SSIV were compared as the Sensi and Omni enzymes failed to produce PCR amplicons for sequencing (as reflected by Ct values above 30; Figure 3). The combination of Sensi and RH priming also failed to reliably amplify *16S rRNA* transcripts and was therefore also excluded from further analysis.

*Effect on 16S rRNA community composition.* When all four enzymes were taken into account, the effect of enzyme on OTU community composition was always significant (Table 4 and Figure 6). In addition, the priming strategy had a significant effect on community composition but only when the Bray-Curtis dissimilarity matrix was considered (Table 4 and Figure 6). Still, the choice of priming strategy had less of an effect on *16S rRNA* community composition than for *amoA,* (Figure 6 and Figure S.3) as GS priming did not systematically result in more OTUs (Figure 7). For the *16S rRNA* dataset, the combined effect of enzyme and priming depended on the specific combination. Specifically, for SSIII and SSIV, there was no difference between enzymes, but there was a significant difference in the Bray-Curtis distance matrix due to priming (richness RH>richness GS for SSIII and inversely for SSIV (Figure 6, 7 and Figure S.4)) albeit marginally significant (*p.*value = 0.047). When Sensi and Omni were compared, both enzyme and priming had an effect on community composition (Table 4, Figure S.4).

**Figure 6.**
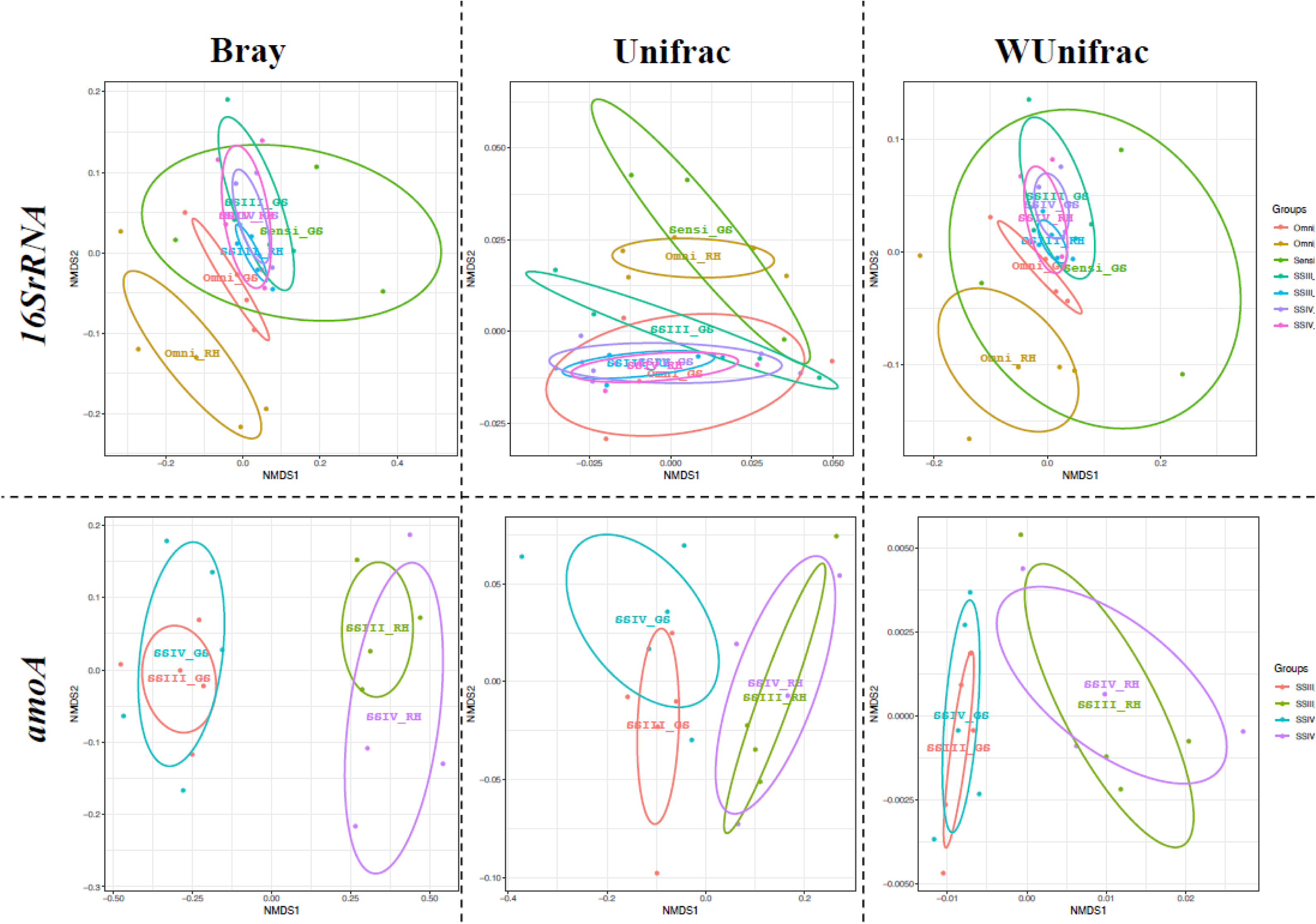
Effect of enzyme and priming on *amoA* and *16S rRNA* transcript community composition. NMDS clustering of *16S rRNA* (top) and *amoA* (bottom) cDNA community composition of the same sample derived from different enzyme and primer strategies, using Bray-Curtis (left), Unifrac (middle) and WUnifrac (right) distances. Corresponding groups are indicated in the legend.

**Figure 7.**
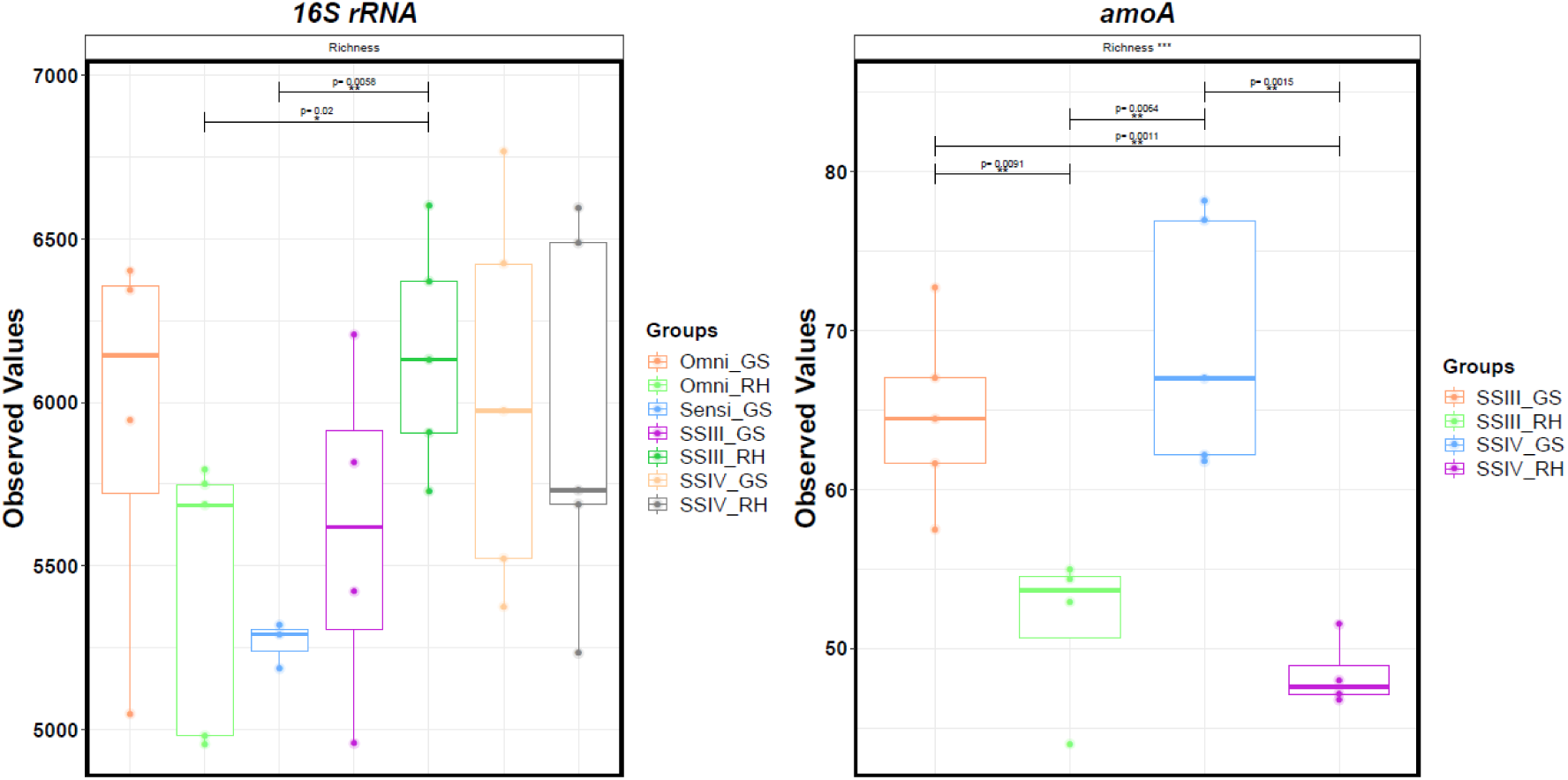
Effect of enzyme and priming on OTU richness. The number of OTUs detected for *16S rRNA* (left) and *amoA* (right) transcripts for the same sample using different RT systems was compared using two-way ANOVA. Results of the statistical tests are represented as lines on top of the plots. *: *p*.value<0.05; **: *p*.value<0.01; ***: *p*.value<0.001.

**Table 4.**
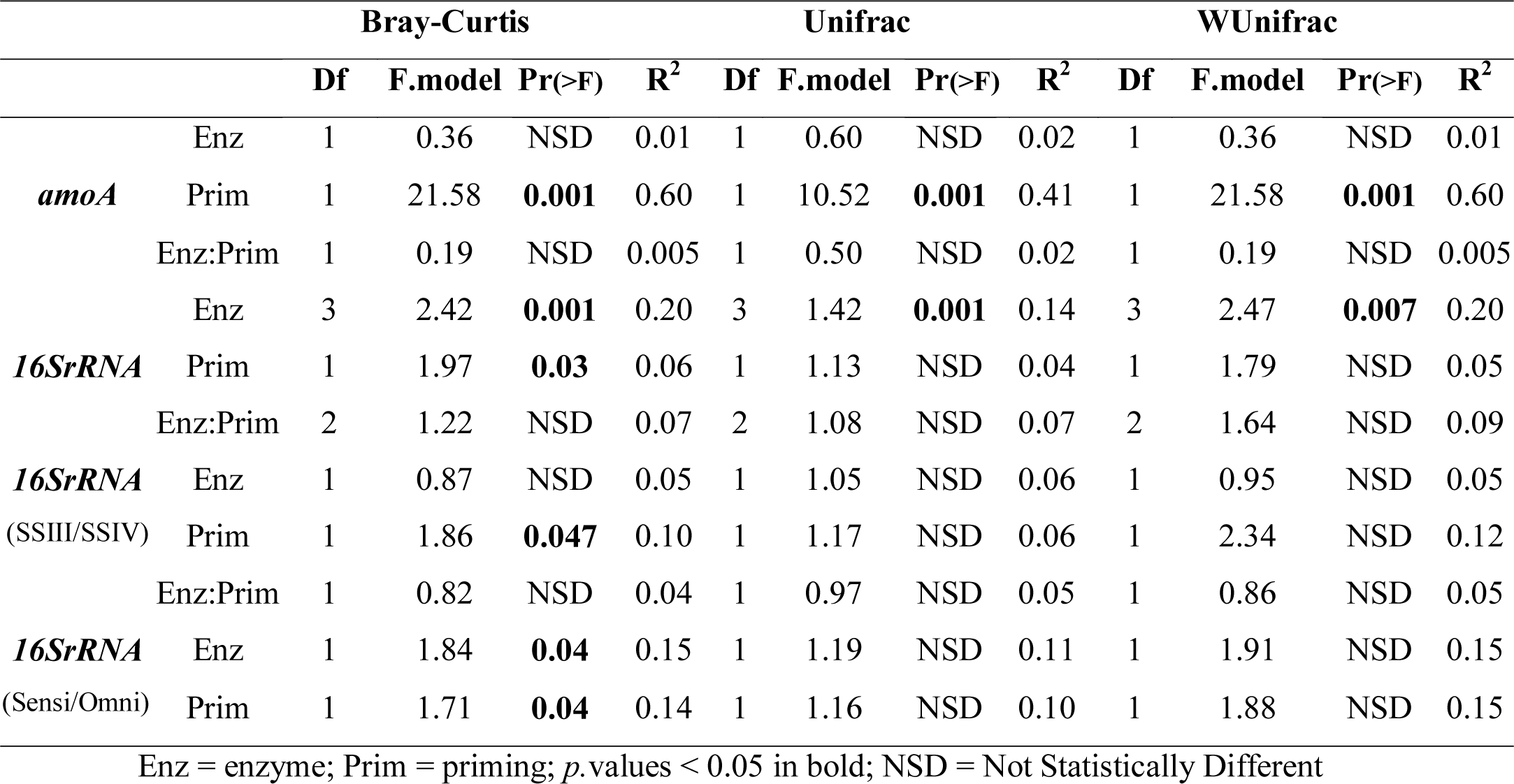
Effect of enzyme and priming on amoA and 16S rRNA community composition: summary of 2-ways PERMANOVA tests.

We further tested the effect of the RT system on the recovery of the *16S rRNA* transcripts at different taxonomic levels (Figure S.3, Table S.4). The effect of enzyme was stronger than priming and was more important when the individual OTUs had well resolved taxonomy (at Family, Order and Class level). On the other hand, at lower taxonomic levels, the effect of the RT system became non-significant as a lot of OTUs could not be assigned to a species or genus and were therefore classified as unknown. Interestingly, the effect of both enzyme and priming became significant again at the kingdom level (Figure S.3, Table S.4).

*amoA OTU check. amoA* OTUs sequences were checked by BLASTx to ensure they translated into AMO A proteins. Results of this search revealed that, out of the 202 *amoA* OTUs, 63 did not correctly translate (*e.g.* “hypothetical protein” or “low quality protein”) and were therefore removed from the data set. In terms of percentage of reads, these non-translating OTUs represented 0.017% to 4.6% of the total. As shown in Figure S.5, the amount of “incorrect OTUs” found was higher in the data set obtained when using GS priming. However, as the number of reads obtained with GS was generally higher, they did not represent a significantly higher percentage of the community, except for the replicate 1 and 3 with SSIV GS where the percentage of incorrect OTUs represented 4.6% and 1.3% respectively (Figure S.5). These OTUs were removed before further processing, and therefore didn’t impact on the subsequent analysis.

*Effect on amoA community composition.* The outcome of *amoA* amplicon sequencing was more strongly influenced by the choice of priming (RH versus GS) than enzyme (SSIII versus SSIV) (Figure 6, Table 4, Figure S.6). In fact, the effect of enzyme on community composition was not significant. On the other hand, priming strategy resulted in a clear, statistically significant clustering of samples (Figure 6, Table 4). One sample, Env1, when prepared using RH priming for both SSIII and SSIV, failed to produce sufficient reads (more than 5000) to proceed and was removed from the analysis pipeline. In contrast, when GS priming was used, sufficient reads were produced to pass this quality step in the analysis pipeline. Indeed, GS priming always resulted in greater OTU richness (Figure 7) than RH (+13 and +21 OTUs on average for SSIII and SSIV respectively), indicating that this priming option was better at recovering the diversity of *amoA* transcripts in the samples. This observation supports the Q-PCR results where GS priming always resulted in lower Cts. To determine if the “missing OTUs” in the RH sequencing data sets were dominant or rare phylotypes, the mean abundance of OTUs present only in the GS data set was plotted for each individual OTU (Figure 8), revealing that most of the OTUs missing in the RH data set were low abundance OTUs. On the other hand, interestingly, a very small number of rare OTUs were only detected in the RH data set (Figure 8). Moreover, the choice of priming also affected the representation of OTUs present in both GS and RH datasets (Figure 8, Table S.5): with SSIII, 39 OTUs were significantly differentially expressed between GS and RH. With SSIV, it was found that 23 OTUs were significantly differentially expressed between GS and RH (Figure 8).

**Figure 8.**
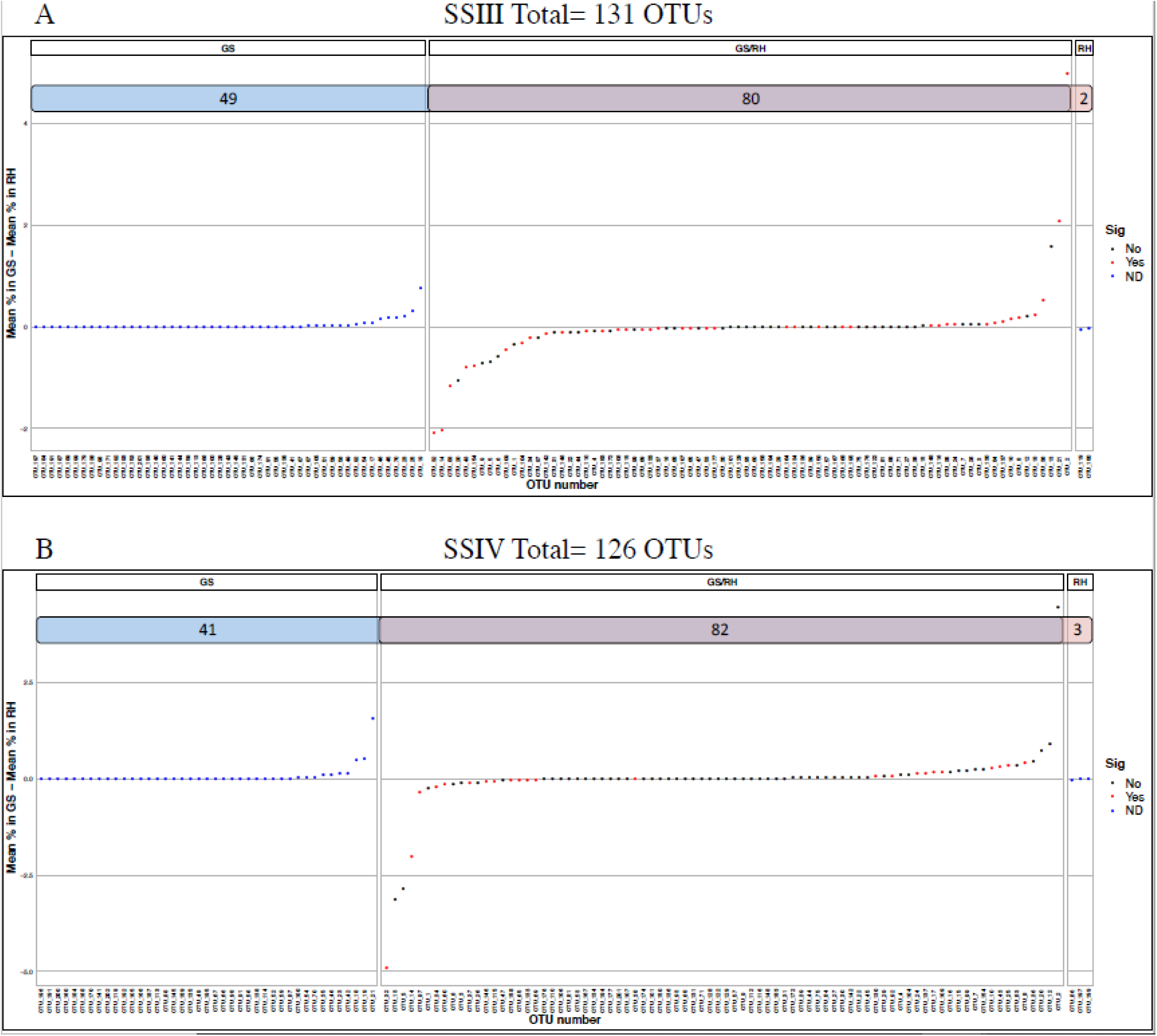
Differences in expression and number of *am* OTUs detected by GS and priming. The Venn Diagrams top of the plots show: the num of OTUs found in the GS data only (blue), in both the GS Rh data set (purple) and in the data set only (red). Results presented as the aver differences in proportion between GS and RH data s (OTUs with positive values overexpressed in the GS inversely). When OTUs w found in only one data set, results are presented as average proportion of the OT with positive and negative va for GS and RH respectively. the OTUs shared between GS RH, the colour of the poi indicates if the difference expression is significant or not explained in the legend (s ND= Not Determined.

## Discussion

While RT-Q-PCR, and to a lesser extent RT-PCR amplicon sequencing, is widely used in environmental microbiology to quantify and determine the diversity of transcripts from environmental samples, the effectiveness and reproducibility of the reverse transcription step has not been evaluated. In particular, in complex environmental samples, to the best of our knowledge, there have not been any studies investigating the efficiency of the reverse transcriptase reaction to transcribe RNA to cDNA, despite this being a critical step informing the overall result. Furthermore, based on our own observations in the laboratory, we often noted the impact of different enzyme and primer choice on the same template. Therefore, we assessed the effect of the RT system (enzyme and priming strategy) on RT-Q-PCR and RT-PCR-amplicon sequencing and showed that the choice of enzyme and priming strategy can result in significant difference in both quantitative and qualitative results from the exact same sample. These methodological effects can bias and even alter final conclusions and interpretations of the underlying biological and ecological questions.

From the *sfGFP* spike experiments (Figures 2 and 3), we showed that the choice of enzyme and priming greatly affected the results of the RT-Q-PCR. When the *sfGFP* transcript was spiked into an environmental RNA background, it was found that the Superscript enzymes performed better than the Sensiscript and Omniscript enzymes. The Superscript enzymes systematically produced higher detected copy numbers, with values closer to the expected ones and, generally, differential expressions closer to the expected 5-fold difference. In a study by Levesque-Sergerie and co-workers (Levesque-Sergerie *et al*., 2007) it was found that the Sensiscript and Omniscript enzymes had a dynamic range >50ng RNA versus >0.01ng RNA for Superscript III. Results obtained here are in accordance, with a better detection of the low concentration target by the Superscript enzymes compared to Sensiscript/Omniscript, especially when RH priming was used. Yet, the RT reactions for standard curves constructed using Sensiscript and Omniscript produced reliable Cts at target concentrations as low as 10^3^ copies/µl, similar to that observed for SSIV and SSIII (except for Sensi-RH: lower limit at 10^4^ copies/µl). This indicates that the lower performances observed for Sensiscript and Omniscript in the environmental spike experiment could be due to inhibition of the enzymes from co-extracted components in environmental RNA (Hata *et al*., 2015) and/or the presence of background RNA. The later explanation contrasts with the results obtained by Levesque-Sergerie and co-workers (Levesque-Sergerie *et al*., 2007) who observed a general increase in the recovered copies of a spike (*i.e.* lower Cts) as the concentration of background RNA from bovine tissue increased.

In this study, GS priming always performed better than RH for RT-Q-PCR, with higher copy numbers and values closer to the expected ones for the exogenous RNA spike. For the endogenous targets (*amoA* and *16S rRNA*) a similar trend was observed, with GS priming resulting in higher detected average copy numbers, except for SSIII, in the *amoA* assay and Omni in the *16S rRNA* assay. The differences between priming were particularly strong with the use of Sensiscript, where the combination of this enzyme and RH was clearly the least efficient RT strategy. Interestingly, small differences were observed between GS and RH when used with SSIV for the quantification of both the spiked *sfGFP* and the endogenous *amoA* showing that this enzyme reliably reverse-transcribed mRNAs. This was also supported by the differential expressions of the exogenous spiked *sfGFP* always being close to the expected 5-fold difference when using SSIV.

Differences in the performances of the RT enzymes and the priming strategies were similar between the *sfGFP* spike and the endogenous *amoA* mRNA with Superscript enzymes performing better than Sensiscript/Omniscript and GS generally performing better that RH. In contrast, for *16S rRNA* significant differences were detected only between Omniscript and the other three enzymes (no statistically significant differences between SSIII, SSIV and Sensiscript) and the effect of priming was very important for all four enzymes. For this assay, Omniscript in combination with RH priming yielded the highest copies/µg RNA. These differences could be a reflection of target concentration, *i.e.* highly abundant *16S rRNA* verses low abundance *amoA* transcripts, or indeed could be target dependant (*i.e.* ribosomal VS mRNA) reflecting for example the complex secondary structure of the RNA molecule.

Overall this study showed that the combination of Superscript IV with GS priming was the most accurate for the quantification of the exogenous *sfGFP* spike and showed the lowest variation in quantification when priming was changed to RH. SSIV GS was also the RT system yielding, on average, the highest copy number for the quantification of *amoA* mRNAs by RT-Q-PCR, coupled with the best precision (lowest standard deviation). This fits with our previous observations, where we would routinely achieve better results (*e.g.* detection verses no detection) and our subsequent choice of the Superscript enzyme with gene specific priming to quantify a range of N-cycle mRNA targets (Duff *et al*., 2017, Smith *et al.,* 2015, 2007).

Next, we investigated the effect of the RT strategy on cDNA sequencing. The result from cDNA sequencing demonstrated that the enzyme and priming strategy employed has an impact on cDNA amplicon diversity. As Sensiscript and Omniscript failed to reliably produce sufficient cDNA to produce PCR amplicons for *amoA*, they were not included, nor was the combination of Sensiscript and RH for the *16S rRNA* diversity study. We have shown that for *amoA* transcripts, priming is an important consideration (Figures 6, 7 and 8; Table 4). Most notably, the use of RH for sample Env1, resulted in too few sequencing reads (<5000) for further analysis. We attribute this to the lower abundance of the *amoA* transcript in a high background of RNA. In this case, the choice of priming made the difference between the success or not of the amplicon sequencing of the transcript. This result was in line with observations from the RT-Q-PCR for *amoA* (Figure 4). Overall more OTUs (Figures 7 and 8) and better coverage of *amoA* transcript diversity (Figure S.7) were obtained when GS priming was used. The differences in the number of OTUs detected was particularly important for low-abundance OTUs indicating that GS priming was better for the reverse transcription of rare members of the *amoA* community (Figure 8). A possible explanation is that, for GS priming, all the RT resources (enzyme and dNTPs) are directed to the reverse transcription of the target transcript. On the other hand, when using RH priming, random priming may not be sufficient to prime rare mRNA target.

This observation is further supported by the results from the *16S rRNA* assay, where the choice of priming strategy was seen to be less important. In fact, here most of the differences observed were due to enzyme choice and not priming strategy. In contrast to the *amoA* results, for *16S rRNA* the use of GS priming did not necessarily result in a higher number of OTUs compared to RH even though GS priming resulted in a higher number of *16S rRNA* copies detected by Q-PCR. It may be that differences in RT performances are abundance or target molecule dependant (*i.e.* very abundant ribosomal RNA with complex secondary structures versus rare messenger RNA).

As the true representation of our transcripts in the environmental samples was unknown, we tested the RT systems against artificial defined RNA mock communities seeded into background environmental RNA. These artificial sequences were derived from target inserts with additional cloning vector sequence added, which allowed for their selective amplification from the background. To evaluate the bias introduced by the PCR/sequencing steps and separate them from the RT, a similar experiment was carried out using DNA mock communities. This experiment revealed that, for all RT systems, biases were introduced in both the RT and the subsequent PCR step of the reaction as the recovered proportions deviated from the expected ones (Figure 5; Figure S.2). When testing a new approach for *16S rRNA* transcript sequencing based on ligation of an adapter to the end of the gene prior to RT with random hexamers, Yan *et al*., 2017, found errors in the observed ratios of their RNA mock communities of up to 3-fold compared to the expected proportions. These results are comparable to those found in this study. Here, we found that the smallest amount of variation from the expected EM composition was observed with SSIV RH. In fact, surprisingly, RH priming always conserved the actual proportions better than GS priming in the seeded mock communities, as seen by lower standard deviations (Figure S.2). Considering that the RNA template for the mock community construction went through both *in vitro*-transcription and a RT reaction prior to PCR, each of which could introduce errors, the standard deviation observed in the RNA mock communities (*i.e.* both GS and RH) was low and in fact, for RH, the same as the DNA mock (4.97 for SSIII GS; 3.31 for SSIII RH; 3.89 for SSIV GS; 3.15 for SSIV RH and 3.34 for DNA) (Figure S.2).

As anticipated, errors were also seen in the UM resulting in observed regressions deviating from the expected. Interestingly, the errors were consistent between EM and UM (*i.e.* a sequence over-represented in the EM would also be over-represented in the UM and vice versa). As a result, when the UM proportions were corrected with the EM ones, the observed regressions were close to the expected y=x (Figure 5). Since the mock communities were constructed separately (Figure 1), this indicated that: 1) these errors are a reflection of sequence specific bias of the RT-PCR workflow and not attributed to user error such as pipetting; 2) Since artificial over/under representations is likely introduced by sequence specific bias, the relative abundance of transcripts within a sample (α diversity) might not always be absolute when small differences (*e.g.* ≈ 4-fold as in this study) in expression are observed; 3) However, as these biases are reproducible (UM reads corrected by EM reads), comparison between samples (*i.e.* β diversity) can be undertaken.

In a recent review about the use of RT-Q-PCR, Bustin and Nolan (Bustin & Nolan, 2017) stated that *“the majority of published RT-Q-PCR data are likely to represent technical noise”*. The intrinsic variability of the RT step and the lack of information on protocols used were key points that lead them to this striking conclusion. This is likely to be similar, if not further amplified in complex environmental samples, from which ecosystem conclusions are drawn. Here we have shown that primer and RT system choice can range from no detection to a 600-fold difference in transcripts for the same template. In environmental studies, this is the difference between no gene expression to the presence of a highly active transcript – striking difference leading to opposite ecosystem conclusions. There is therefore an urgent need to ensure that the approaches we use are tested and recommendations as far as possible for best practice are made, followed and reported in future studies. Our study shows that the choice of correct enzyme and priming can improve the reliability and reproducibility of RT-Q-PCR and RT-sequencing data, facilitating insight into the transcriptionally active microbial communities directly from the environment. This, taken together with steps to monitor the purity and integrity of the extracted RNA prior to downstream analysis (Bustin and Nolan, 2017; Cholet *et al*., 2019) and detailed documentation of the RT approach used should greatly improve the reliability and reproducibility of transcript based studies in environmental microbiology. From our work, we put forward the following recommendations for best practice:

### 1: Evaluate and report RNA quality and integrity

As previously reported (Cholet *et al*., 2019), the quality and in particular, the integrity of the extracted environmental RNA should be determined and reported as the mandatory first step in any RNA based workflow.

### 2: RT-Q-PCR

i. Gene specific priming was more accurate, precise and sensitive than random hexamer priming for mRNA.
ii. Of the enzymes tested, Superscript IV was accurate, precise and sensitive, and therefore we recommend its use for the detection of transcripts in complex environmental RNA matrixes.
iii. The incorporation of an exogenous RNA target at known concentration into the environmental RNA being tested is an efficient way to validate RT-Q-PCR protocols.
iv. When converting Ct results into copy number, we advise the use of an RNA standard curve (*i.e.* serial dilution of the target RNA and individual RT-Q-PCR) rather than a cDNA standard curve (*i.e.* reverse transcription of a fixed concentration of RNA, dilution of the cDNA and Q-PCR).
v. Fully report the RT protocol used.

### 3: RT-amplicon sequencing

i. For RT-amplicon sequencing of mRNA targets, we recommend the use of gene specific priming as it resulted in better coverage and higher OTU richness of the bacterial *amoA* transcript. For *16S rRNA* RT-sequencing, the choice of priming is less important.
ii. The addition of RNA mock communities into environmental RNA (before reverse transcription) can aid interpret sequencing results: in our case, we deduced from our RNA mock communities that even though relative proportions of individual OTUs within a sample (α diversity) can be biased, the comparisons of changes in OTU composition between samples (β diversity) are reliable.

### Experimental Procedures

#### Sediment sample collection

Surface mud samples (0 to 2 cm) were collected on 11/01/2017 from Rusheen Bay, Ireland (53.2589° N, 9.1203° W) (presence of *amoA* genes/transcripts previously established (Duff *et al*., 2017; Zhang *et al*., 2018, Cholet *et al.,* 2019) in sterile 50ml Eppendorf tubes, flash frozen and stored at −80°C until subsequent use. Five biological replicate sediments, designated Env1, Env2, Env3, Env4 and Env5 respectively were used for testing the effect of the RT reaction on RT-Q-PCR and RT-amplicon sequencing of the endogenous *amoA* and *16S rRNA* transcripts. An additional sample was used for preparing the RNA background for the *sfGFP* spiking experiment.

#### RNA preparation from sediment

All surfaces and equipment were cleaned with 70% ethanol and RNase Zap (Ambion) before sample processing. All glassware and stirrers used for solutions were baked at 180°C overnight to inactivate RNases. All plasticware was soaked overnight in RNase away solution (ThermoFisher Scientific). Consumables used, including tubes and pipette tips were RNase free. All solutions were prepared using Diethylpyrocarbonate (DEPC) treated Milli-Q water. A simultaneous DNA/RNA extraction method, based on that of Griffiths and co-workers (Griffiths *et al*., 2000) was used to recover nucleic acids from sediment. Briefly, 0.5g of sediments were extracted using bead beating lysing tubes (Matrix tube E; MP Biomedical) and homogeneised in 0.5ml CTAB/phosphate buffer (composition for 120 ml: 2.58g K_2_HPO_4_.3H_2_O; 0.10g KH_2_PO_4_; 5.0g CTAB; 2.05g NaCl) plus 0.5ml

Phenol:Chlorophorm:Isoamyl alcohol (25:24:1 v:v:v). Lysis was carried out on the FastPrep system (MP Biomedical) (S: 6.0; 40sec) followed by a centrifugation at 12,000g for 20 min 4°C). The top aqueous layer was transferred to a fresh 1.5ml tube and mixed with 0.5ml chloroform:isoamyl alcohol (24:1 v:v). The mixture was centrifuged at 16,000g for 5 min (4°C) and the top aqueous layer was transferred to a new 1.5ml tube. Nucleic acids were precipitated by adding two volumes of a solution containing 30% poly(etlyleneglycol)_6000_ (PEG6000) and 1.6M NaCl for 2 hours on ice and subsequently recovered by centrifugation at 16,000 x g for 30 min (4°C). The pellet was washed with 1ml ice-cold 70% ethanol and centrifuged at 16,000g for 30 min (4°C). The ethanol wash was discarded, and the pellet was air dried and re-suspended in 40µl DEPC treated water. DNA/RNA preparations were stored at −80°C if not used immediately. RNA was prepared from the DNA/RNA co-extraction by DNase treating with Turbo DNase Kit (Ambion) using the extended protocol: half the recommended DNase volume is added to the sample and incubated for 30 min at 37°C, after which the second half of DNase is added, and the sample is re-incubated at 37°C for 1 hour. Success of the DNase treatment was checked by no PCR amplification of the V1-V3 Bacterial *16S rRNA* gene (Smith *et al*., 2006).

### RNA quality check

The quality, purity and integrity of extracted environmental RNA was determined as follows: *Quantity/purity*: Total RNA was quantified using three different approaches: spectrophotometry (NanoDrop; Life Technologies), fluorometry (Qubit broad Range RNA; Life Technologies) and microfluidics (Bioanalyser 2100 RNA Nano; Agilent Technologies). Purity was determined by spectrophotometry (NanoDrop; Life Technologies) with the 260nm/230nm and 260nm/280nm band absorption ratios.

*Integrity*: RNA integrity was determined using two different approaches: the RNA Integrity Number (RIN), based on the 23S/16S rRNA ratio and the electropherogram of the extracted RNA (Bioanalyser 2100 RNA Nano; Agilent Technologies) and the R_amp_ approach, based on the differential amplification of glnA mRNA amplicons of different length (Cholet *et al.,* 2019).

### Evaluation of RT reaction on transcript quantification via a sfGFP RNA spike

#### Preparation of the sfGFP RNA

A plasmid containing the *sfGFP* (pTHSSd_8) gene (designed by Segall-Shapiro et al. (2014)) was ordered from the Addgene plasmid repository web site (https://www.addgene.org/). A DNA preparation of the plasmid was prepared for PCR and Q-PCR as the target to optimise primers, initially at DNA level and subsequently for RT-Q-PCR as appropriate. The amplicon obtained by PCR amplification using pBRforECO and GFP-Frc (Table S.6) was then used as template for *in vitro* transcription to produce the RNA spike (see Supplementary Experimental procedure for details).

#### sfGFP Q-PCR standard curves

*sfGFP* RNA dilutions (10^10^ 10^1^ transcript copies/µl) were prepared and individually reverse transcribed (RT) using four different enzymes: Superscript IV (SSIV), Superscript III (SSIII) (Invitrogen), Sensiscript (Sensi) and Omniscript (Omni) (Qiagen) and two priming strategies - gene specific (GS) and random hexamer (RH). Each RT was done in duplicate. A summary of the protocol for each system is presented in Table S.7. The resulting cDNA preparations underwent Q-PCR using the primer pair sF300_Fand sF300_R (Table S.6). Each 20µl Q-PCR reaction contained 10µl EVAGreen Supermixes 2X (SsoFast; Bio-Rad), 0.5µl of each primer (10µM each), 8µl water and 1µl of cDNA template. Further details are provided in the Supplementary Information file.

#### Spiking experiment

In order to determine which enzyme/priming combination was the most accurate, the exogenous RNA spike (*sfGFP* RNA) was seeded into a background of environmental RNA ([RNA]_background_ =70.7 ng/µl; ratio 260/280_background_ = 1.63; ratio 260/230_background_ = 0.87) at known concentrations: 10^3^, 5 x 10^3^, 2 x 10^6^ and 10^7^ copies/µl. The RNA background was same for all spikes. These concentrations were chosen to mimic five-fold changes in gene expression at both low and high expression level. After the *sfGFP* spike was added, total RNA was reverse transcribed in triplicate, using different combinations of enzymes and priming (four different RT enzymes; two different priming strategies) in the same manner as illustrated in Figure 1 and Table S.7. A 300bp fragment of the *sfGFP* cDNA was then quantified from the cDNA preparations using quantitative PCR (one Q-PCR reaction for each of the 3 RT replicates) with the primer pair sF300_F/ sF300_R. The Q-PCR mix was composed of 10µl EVAGreen Supermixes 2X (SsoFast; Bio-Rad), 0.5µl each primer (10µM each), 8µl water and 1µl cDNA template.

### Differential Expression (DE) between consecutive spike concentrations

The fold difference between consecutive spike concentrations was then calculated as the ratio of the mean copies/µl exogenous spike detected: DE “Low” corresponds to the ratio of mean copies/µl detected in the 10^3^ spike versus the 5 x 10^3^ spike. DE “High” corresponds to the ratio of mean copies/µl detected in the 2 x 10^6^ spike versus the 10^7^ spike. The standard deviations of the ratios were calculated as:

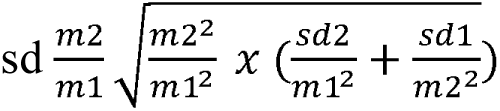

where m1 is the mean copies/µl at concentration C and m2 the mean copies/µl at concentration C x 5; sd1 and sd2 the standard deviations of m1 and m2 respectively.

### Effect of RT reaction on quantification of endogenous amoA and 16S rRNA transcripts

#### amoA and 16S rRNA Q-PCR standard curves

*<u>amoA and 16S rRNA</u>* RNA dilutions were prepared and each individually reverse transcribed (RT) using four different enzymes: Superscript IV (SSIV), Superscript III (SSIII) (Invitrogen), Sensiscript (Sensi) and Omniscript (Omni) (Qiagen) and two priming strategies - gene specific (GS) and random hexamer (RH). A summary of the protocol for each system is presented in Table S.7. The resulting cDNA preparations underwent Q-PCR using the corresponding primer pair and Q-PCR conditions as detailed in Table S.6 (regression coefficients in Table S.6).

#### RT-Q-PCR endogenous transcripts

RNA was extracted from five biological replicates (marine sediment samples) and reverse transcribed as described in Figure 1 and Table S.7. The *amoA* Q-PCR was carried out in 20µl reaction volume composed of 10µl EVA Green master mix, 0.4µl of each primer (BacamoA-1F and BacamoA-2R) (10µM each), 7.2µl water and 2µl of cDNA template (1/10 diluted). *16S rRNA* cDNA targets were quantified in a 20µl reaction volume composed of 10µl Itaq Universal Probes Supermix (Bio-Rad), 1.8µl each primer (1369F and 1492r) (10µM each), 0.4µl probe (1389P) (10µM), 5µl water and 1µl cDNA template (1/10 diluted).

### Evaluation of RT reaction on transcript community composition

#### Effect of reverse transcription on sequencing of endogenous transcripts

Besides evaluating the effect of the RT enzyme and priming strategy on the quantification of the endogenous *amoA* and *16S rRNA* transcripts, the effect of these on community composition, as determined by amplicon sequencing of cDNA was also studied. To this end, *amoA* and *16Sr RNA* amplicons were generated from the cDNA preparations used in the previous experiment (Figure 1.B). For the *amoA* transcript, only SSIII and SSIV enzymes with both random hexamer and gene specific priming strategy were considered, as the other enzyme systems failed to work. For *16S rRNA* all four enzymes and both priming strategies produced amplicons and were therefore tested. Details for MiSeq-Illumina amplicon library preparation are provided in the section “MiSeq Illumina sequencing”.

### Effect of reverse transcription on sequencing of exogenous tagged-mock community

#### Preparation of the mock communities

*16S rRNA* amplicons were generated by PCR amplification of the V4 region of the Bacterial *16S rRNA* gene from environmental samples using primers 515F and 806R (Table S.6). PCR products were cloned and sequenced from which 12 different sequences (97% similarity threshold) were selected to make a mock community from. For the DNA mock communities, PCR amplicons from the 12 individual clones, were quantified and pooled together in different proportions to create an Even Mock (EM) and an Uneven Mock (UM) communities: for the EM, each sequence was represented at even proportions while for the UM, sequences were pooled at different proportions, following a log-normal distribution (Table S.3). RNA mock communities were constructed by *in-vitro* transcription of the individual 12 PCR amplicons used in DNA mock communities. The 12 individual RNA preparations were quantified and pooled together to obtain the EM and UM as for the DNA mock (Table S.3). A detailed procedure for the construction of the DNA and RNA mock is provided in Supplementary Material and Methods 1.

#### Spiking and recovery of the RNA mock communities

Once constructed, the RNA mock communities (EM and UM) were diluted 1/10 into environmental RNA background (background: [RNA] = 102.3 ng/µl; 260/280 = 1.65; 260/230 = 1.28). This step was repeated five times. Once seeded with the mock communities, the environmental RNA preparations were reverse transcribed using two different enzymes (SSIII and SSIV) and two different priming strategies (RH and GS) (Figure 1 C). The GS priming was carried out using the 806R primer (Table S.6). The procedure followed was the same as described in Table S.7. After reverse transcription, the spiked mocks sequences were recovered from the total RNA pool by PCR amplification using the 806R reverse primer and a custom vector-specific forward primer pGEMT_FW2 (Table S.6), designed to amplifying pGEMT vector sequence located between the T7 promotor site and the beginning of the insert, hence insuring the specific amplification of the mock sequences from the background (Figure 1). The specificity of the pGEMT_FW2 forward primer was checked by the absence of amplification of environmental *16Sr RNA* genes when used in combination with 806R.

### Illumina MiSeq amplicon library preparation

Primers used for sequencing were modified by adding Illumina adaptors at the 5’ end: 5’-TCG TCG GCA GCG TCA GAT GTG TAT AAG AGA CAG (forward adaptor); 5’-GTC TCG TGG GCT CGG AGA TGT GTA TAA GAG ACA G (reverse adaptor). The use of vector-targeting forward primer ensured that only the spiked mock communities were amplified. The specificity of this PCR assay was verified by the absence of amplification from the un-spiked reverse transcribed background. The PCR was carried out using the HotStartTaq PCR kit (Qiagen) in a 25µl volume: 19.8µl water, 0.5µl of each primer (10µM each), 0.5µl dNTPs (10µM each), 0.2µl HotStartTaq, 2.5µl of 10x PCR buffer and 1µl cDNA template (10^-1^). For the *amoA* functional gene, three separate PCRs were carried out per sample and pooled together for further processing. PCR amplicons were cleaned using the Agencourt AMPure XP beads (Beckman Coulter) following the manufacturer’s recommendations. Illumina indexes were attached using the Nextera XT Index Kit with the following PCR condition: 95°C-15min, (95°C-30sec, 55°C-30sec, 72°C-30sec) x 8 cycles and 72°C-5min. The resulting amplicons were purified using the Agencourt AMPure XP beads (Beckman Coulter) and eluted in 25µl water. After this step, some preparations were randomly chosen (two per gene target) and analysed on the Bioanalyser using the DNA 1000 Assay protocol (Agilent Technologies) to determine the average length of the amplicons and to check for the presence of unspecific products. Finally, DNA concentration was determined using fluorometric quantification method (Qubit) and molarity was calculated using the following equation: (concentration in ng/ μl) × 10^6^ = (660 g/mol × average library size).

Libraries were pooled in equimolar amount and checked again on the Bioanalyser and the final library was sent to the Earlham Institute (Norwich Research Park, Norwich, UK) for Illumina MiSeq amplicon sequencing (300PE, 22 millions reads/ lane).

### Processing of amplicon sequences

#### Construction of the reference databases

The following sequences were downloaded (see Additional file 2): *amoA* sequences from FunGene (http://fungene.cme.msu.edu/) alongside NCBI sequences (n=642) as FASTA files. The NCBI taxonomy was given in the FASTA headers. Subsequently, R’s rentrez (Winter, 2017) package was used to get taxonomic information at different levels to generate a taxonomy file. The FASTA file and the corresponding taxonomy file was then formatted to work with Qiime. For *16S rRNA* we used the SILVA SSU Ref NR database release v123. For more information, see Cholet *et al.,* 2019.

### Bioinformatics pipeline

Abundance tables were obtained by constructing operational taxonomic units (OTUs) as follows. Paired-end reads were trimmed and filtered using Sickle v1.2 (Joshi & Sickle, 2011) by applying a sliding window approach and trimming regions where the average base quality drops below 20. Following this we apply a 10 bp length threshold to discard reads that fall below this length. We then used BayesHammer (Joshi & Sickle, 2011) from the Spades v2.5.0 assembler to error correct the paired-end reads followed by pandaseq v(2.4) with a minimum overlap of 20 bp to assemble the forward and reverse reads into a single sequence. The above choice of software was as a result of author’s recent work (Schirmer *et al*., 2015; D’Amore *et al*., 2016) where it was shown that the above strategy of read trimming followed by error correction and overlapping reads reduces the substitution rates significantly. After having obtained the consensus sequences from each sample, the VSEARCH (v2.3.4) pipeline (all these steps are documented in https://github.com/torognes/vsearch/wiki/VSEARCH-pipeline) was used for OTU construction. The approach is as follows: the reads are pooled from different samples together and barcodes added to keep an account of the samples these reads originate from. Reads are then de-replicated and sorted by decreasing abundance and singletons discarded. In the next step, the reads are clustered based on 97% similarity, followed by removing clusters that have chimeric models built from more abundant reads (--uchime_denovo option in vsearch). A few chimeras may be missed, especially if they have parents that are absent from the reads or are present with very low abundance. Therefore, in the next step, we use a reference-based chimera filtering step (--uchime_ref option in vsearch) using a gold database (https://www.mothur.org/w/images/f/f1/Silva.gold.bacteria.zip) for *16S rRNA* sequences, and the above created reference databases for *amoA* genes. The original barcoded reads were matched against clean OTUs with 97% similarity to generate OTU tables. The representative OTUs were then taxonomically classified using assign_taxonomy.py script from Qiime (Caporaso *et al*., 2010) against the reference databases. To find the phylogenetic distances between OTUs, we first multi sequence aligned the OTUs against each other using Mafft (Katoh *et al*., 2009) and then used FastTree v2.1.7 (Price *et al*., 2010) to generate the phylogenetic tree in NEWICK format. Finally, make_otu_table.py from Qiime workflow was employed to combine abundance table with taxonomy information to generate biome file for OTUs.

#### amoA OTUs check

To ensure all *amoA* OTUs were valid *amoA* sequences they were translated using BLASTx to proteins and the match recorded for each individual OTU. Results of this search were used to filter the OTU_table before further processing, and non-translated amplicons removed from further analysis.

### Statistical analysis

All statistical analyses were carried out in R (R Core team 2013). The effect of enzyme and priming on RT-Q-PCR result were tested after a log10 transformation of copy number data for 2-way ANOVA tests because the assumption of homogeneity of variances between groups was violated when using copy number directly. When the two-way ANOVA was significant, differences between enzymes/priming strategy were investigated using Tuckey HSD post-hoc test.

## Acknowledgements

FC was supported by a Univeristy of Glasgow, College of Science and Engineering Doctoral Scholarship; UZI was funded by NERC IRF NE/L011956/1; CJS was supported by the Royal Academy of Engineering-Scottish Water Research Chair (RCSRF1718643). We would like to thank Dr. Aoife Duff for collecting sediment samples.

## Data Availability

The sequencing data are available on the European Nucleotide Archive under the study accession number: PRJEB32314 (http://www.ebi.ac.uk/ena/data/view/PRJEB32314) with the details given in sequencing_data_information.xls

## Competing interest

The authors declere that they have no competing interest

## Supplementary Experimental Procedures

### Preparation of the *sfGFP* RNA spike and standard curves

A plasmid containing the *sfGFP* gene (designed by Segall-Shapiro et al. (2014)) was ordered from the Addgene plasmid repository web site (https://www.addgene.org/59948/) as a bacterial stab (*E. coli*). The bacterial stab was streaked on LB agar + Ampicillin (100 µg/ml) and incubated overnight at 37°C. A single colony was re-grown in LB ampicillin (100 µg/ml) and used to generate glycerol stocks and subsequently used for PCR and Q-PCR validation of the primers (see main document). For all PCR amplifications, three negative control were included: *E. coli* DNA, environmental DNA and a no template control. The primer sF500_R (used for gene specific reverse transcription) was also tested at PCR level to ensure its specificity for subsequent RT experiments. After validation of the primers, a fragment of the plasmid including the T7 promotor site and the quasi full length *sfGFP* gene (40 bp at the 3’ end was not included) was PCR amplified using primers pBRforEco and GFP-Frc (reverse complement of GFP-F) (Table 1). The PCR product was purified using Agencourt AMPure XP beads (Beckman Coulter) following the manufacturer’s recommendations and then used for *in vitro* transcription using the MEGAscript T7 transcription kit (Invitrogen) to prepare *sfGFP* RNA. The RNA preparation was treated with the Turbo DNase kit (Ambion) and the full digestion of the DNA template was confirmed by the absence of PCR amplification of a 300bp *sfGPF* fragment using the sF300_F and sF300_R primer pair (Table 1). Production of target RNA of the correct length was confirmed on the 2100 Bioanalyser RNA nano (Agilent) and concentration determined using fluorometric quantification method (Qubit RNA BR assay; ThermoFisher Scientific). The number of RNA transcripts was calculated using EndMemo RNA copy number Calculator (http://endmemo.com/bio/dnacopynum.php). A 10 points serial dilution was prepared by successive 1/10 dilutions. RNA dilutions were reverse transcribed using the combination of four enzymes and two priming (Table S.5.) and the resulting cDNA was diluted 1/10 before Q-PCR amplification

### Preparation of *amoA* and *16SrRNA* RNA standard curves

RNA standard curves were constructed by serial dilution of the target RNA, reverse transcription of the individual dilutions and Q-PCR amplification of the resulting cDNA: First, the genes of interest (Bacterial *amoA* and *16S rRNA*; Table S.4.) were amplified and cloned into the pGem-T-Easy Vector Systems (Promega). The resulting ligation was transformed into the competent cell *E.coli* JM109 by heat shock according to manufacturer’s instruction. White colonies were then PCR screened using T7 (5’-TAATACGACTCACTATAGGG–3’) forward and gene specific reverse primer to ensure the amplicon had been cloned in the 5’ 3’ orientation.

Colonies that gave positive results were re-amplified using M13F (5’-GTAAAACGACGGCCAGT-3’) and M13R (5’-CAGGAAACAGCTATGAC-3’) and the resulting PCR products were purified using the SureClean Plus DNA purification kit (Bioline), then quantified using Qubit DNA High Sensitivity (ThermoFisher Scientific) and used as template for *in-vitro* transcription using the MEGAscript T7 transcription kit (ThermoFisher Scientific). RNA was then purified using the RNA clean-up protocol of the RNeasy Mini Kit (Qiagen) and DNase treated using an extended protocol: for 20µl purified RNA, 1µl DNase was added and incubated for 1h at 37°C. Then, another 1µl DNase was added and further incubated for 1h at 37°C. RNA was recovered using the phenol-chlorophorm/Isopropanol method (as recommended by the MEGAscript instruction manual) and complete digestion of the DNA template was confirmed by no amplification of the insert using T7F and M13R primers. The concentration and size of the DNA-free RNA were then checked using the Bioanalyzer 6000 RNA Nano (Agilent) assay. The number of RNA transcripts was calculated using EndMemo RNA copy number Calculator (http://endmemo.com/bio/dnacopynum.php). For *amoA*, an eight points serial dilution (5^10^ to 5^3^ copies/µl) was prepared by successive 1/5 dilutions. For *16SrRNA*, a five points serial dilution (10^9^ to 10^5^ copies/µl) was prepared by successive 1/10 dilutions. RNA dilutions were reverse transcribed using the combination of four enzymes and two priming (Table S.5.) and the resulting cDNA was diluted 1/10 before Q-PCR amplification (Table S.4. for protocols; Table S.6. for results).

### Preparation of the mock communities

*16S rRNA* amplicons were generated by PCR amplification of the V4 region of the Bacterial *16S rRNA* gene from environmental samples using primers 515F and 806R (Table 1). PCR products were purified using the SureClean Plus kit (Bioline) and ligated into the pGEM-T Easy Vector using the pGEM-T Easy Vector System I (Promega) following manufacturer’s instructions. The resulting constructions were transformed into *E.coli* MDS42 competent cells using the heat shock method (50µl *E.coli* MDS42 competent cells culture was incubated with 2µl of the ligation reaction on ice for 20min and transferred at 42°C for 50sec, then back on ice for 2min). The transformation solution was then added to 950µl SOC medium and incubated at 37°C for 1 hour. 100µl was then plated on LB agar/ampicillin (100µg/ml)/Xgal (20µg/ml)/IPTG (0.5 mM). After overnight incubation at 37°C, white colonies were picked up from the plates and grown for 2h30min in LB ampicillin (100µg/ml). A colony PCR was then carried using the BiotaqRed kit (Bioline) in a 50µl final reaction composed of 25µl 2X BiotaqRed Buffer, 1µl T7 forward primer (10µM), 1µl 806 reverse primer (10µM) and 1µl Bacterial culture (the T7 forward primer was used instead of the 515 forward to ensure that the amplicon had been cloned in the 5’ 3’ orientation relative to the T7 promoter). The rest of the culture was stored as glycerol stock at −80°C. When band of the expected size were detected on agarose gels, the PCR reaction was purified using the SureClean Plus kit (Bioline) and send for Sanger sequencing. In total 47 purified PCR products were sent for Sanger sequencing.

The sequences obtained were submitted to BLAST search (2) and then clustered at 97% similarity. A phylogenetic tree was constructed using MEGA7 (3) and the twelve most dissimilar sequences were selected for the mock community construction. Colony PCRs were then carried out using the glycerol stock of these 12 clones as template. The amplicons were generated using M13 forward and M13 reverse primers (Table 1). The resulting amplicons were composed of the full length insert (≈290bp) plus a vector sequence of 100bp at the 5’ end containing the T7 promoter site, and 137bp at the 3’ end. After purification, the twelve PCR products were quantified using fluorometric quantification method (Qubit DNA High Sensitivity assay) and the corresponding copy number was calculated using EndMemo DNA copy number Calculator (http://endmemo.com/bio/dnacopynum.php). Finally, they were pooled together as explained in Figure 1 and Table S.3 to generate the DNA Even (EM) and Uneven (UM) mock communities. Once pooled together, the DNA mock communities were diluted in order to reach 10^8^ copies/µl for each sequence.

The RNA mock communities were constructed by *in-vitro* transcription of the individual 12 PCR products (from DNA mock) using the MEGAscript T7 transcription kit (Invitrogen). The resulting RNAs were then gel-purified and DNase treated using the Turbo DNase Kit (Ambion). Complete digestion of the DNA template was confirmed by the absence of PCR amplification (using 515F and 806R primers; Table 1). The 12 individual RNA preparations were quantified using fluorometric quantification method (Qubit RNA broad range assay) and the corresponding copy number was calculated using EndMemo RNA copy number Calculator (http://endmemo.com/bio/dnacopynum.php). Each RNA preparation was diluted to 10^10^ copies/µl. Even (EM) and Uneven (UM) RNA mock communities were prepared as explained in Table 3 and Figure 1. Once pooled together, the RNA mock communities were diluted to reach 10^8^ copies/µl for each sequence.

## Supplementary Figures

**Figure S.1.**
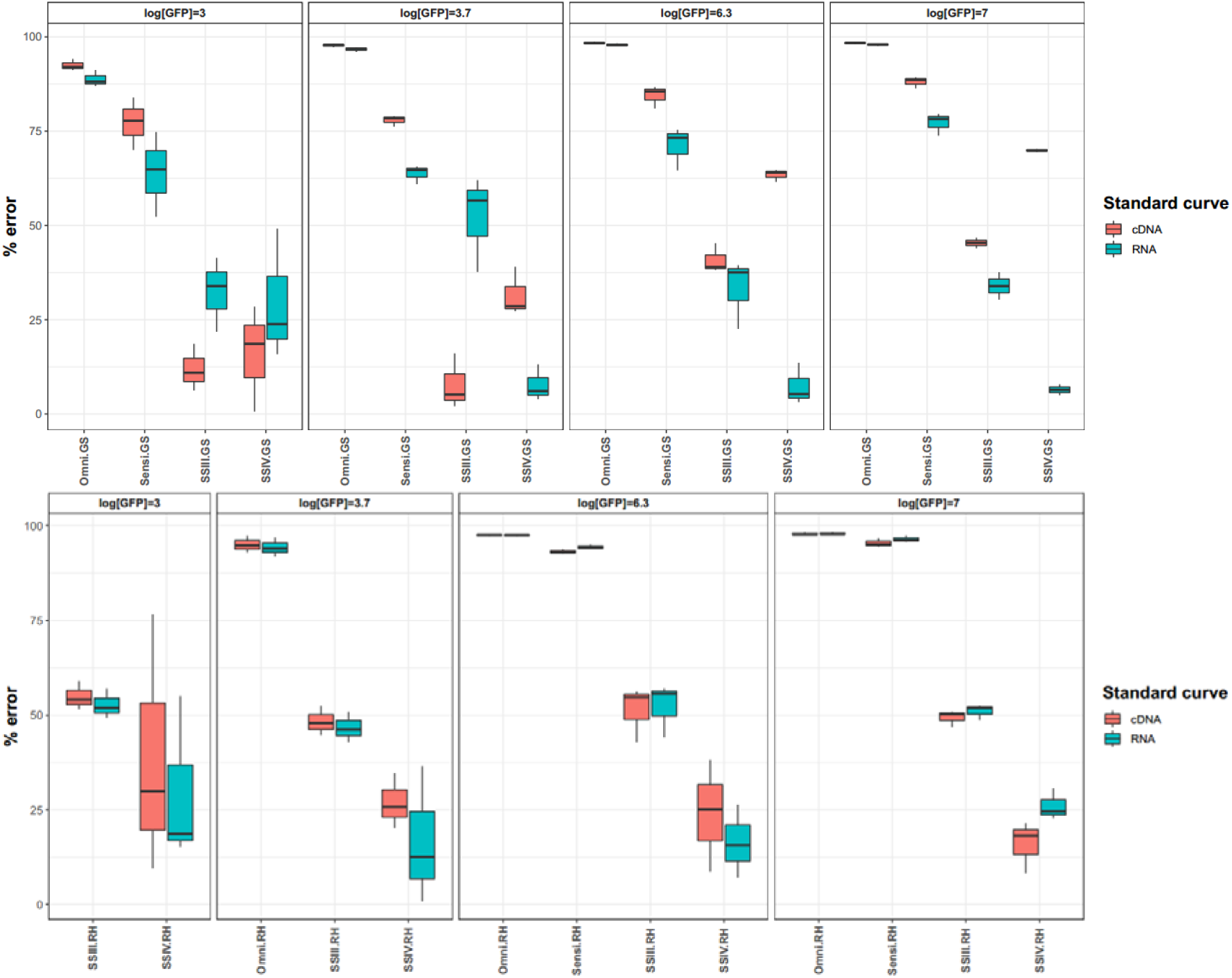
Percentage error in the detected copies/µl of spike *sfGFP*. The percentage error has been calculated for the three replicates and expressed in absolute value regardless of the type of error (over or under estimation of the expected value), for GS (top) and RH (bottom) priming. Standard curve RNA: standard curve made by serial dilution of RNA and individual RT, Standard curve cDNA: standard curve made by a single RT and serial dilution of cDNA.

**Figure S.2.**
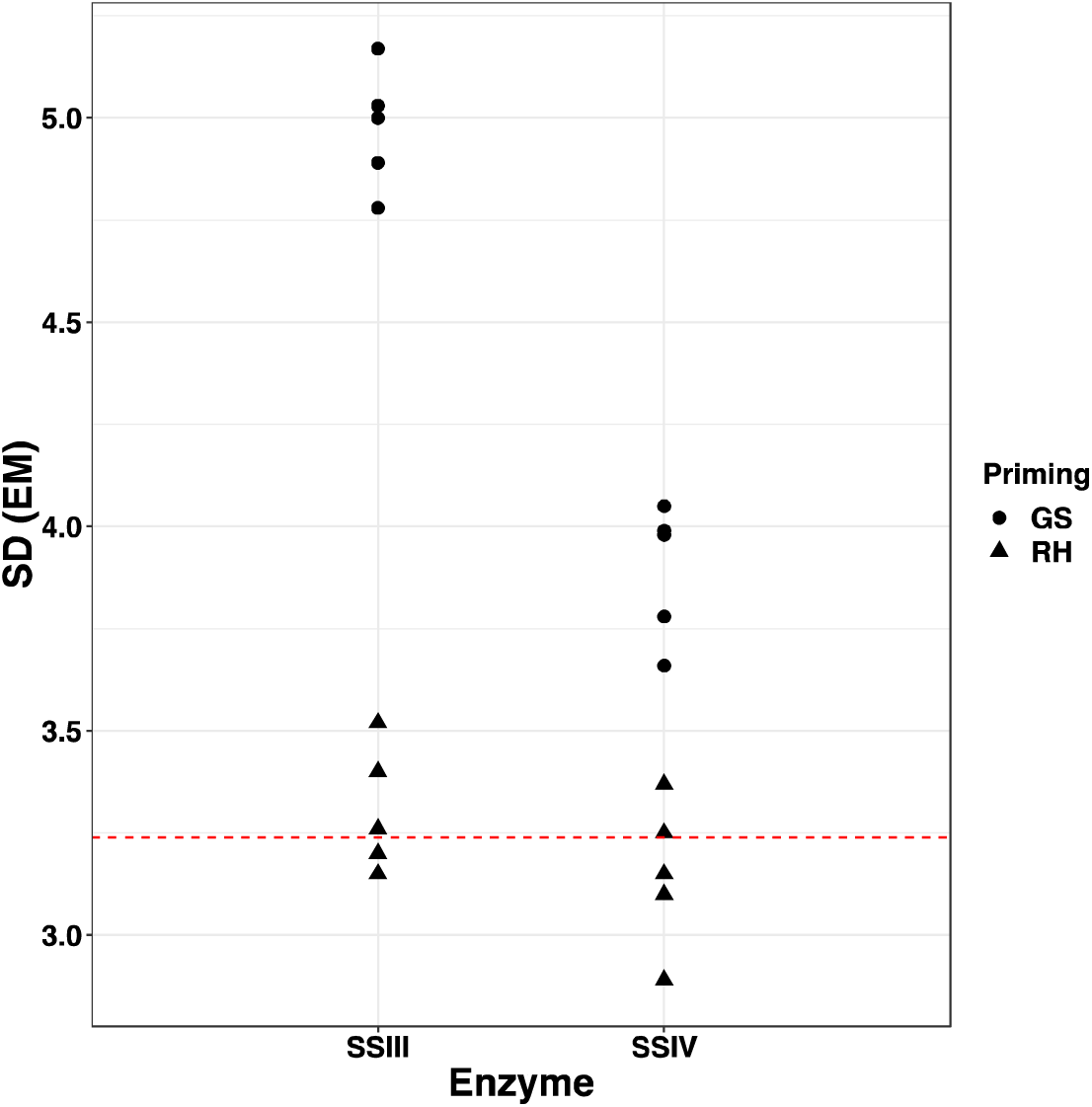
Errors in the sequencing of the cDNA even mocks. The errors are represented as the standard deviations (sd) of the 5 EM replicates. For reference, the mean sd observed in the DNA mock is indicated as a red dashed line

**Figure S.3.**
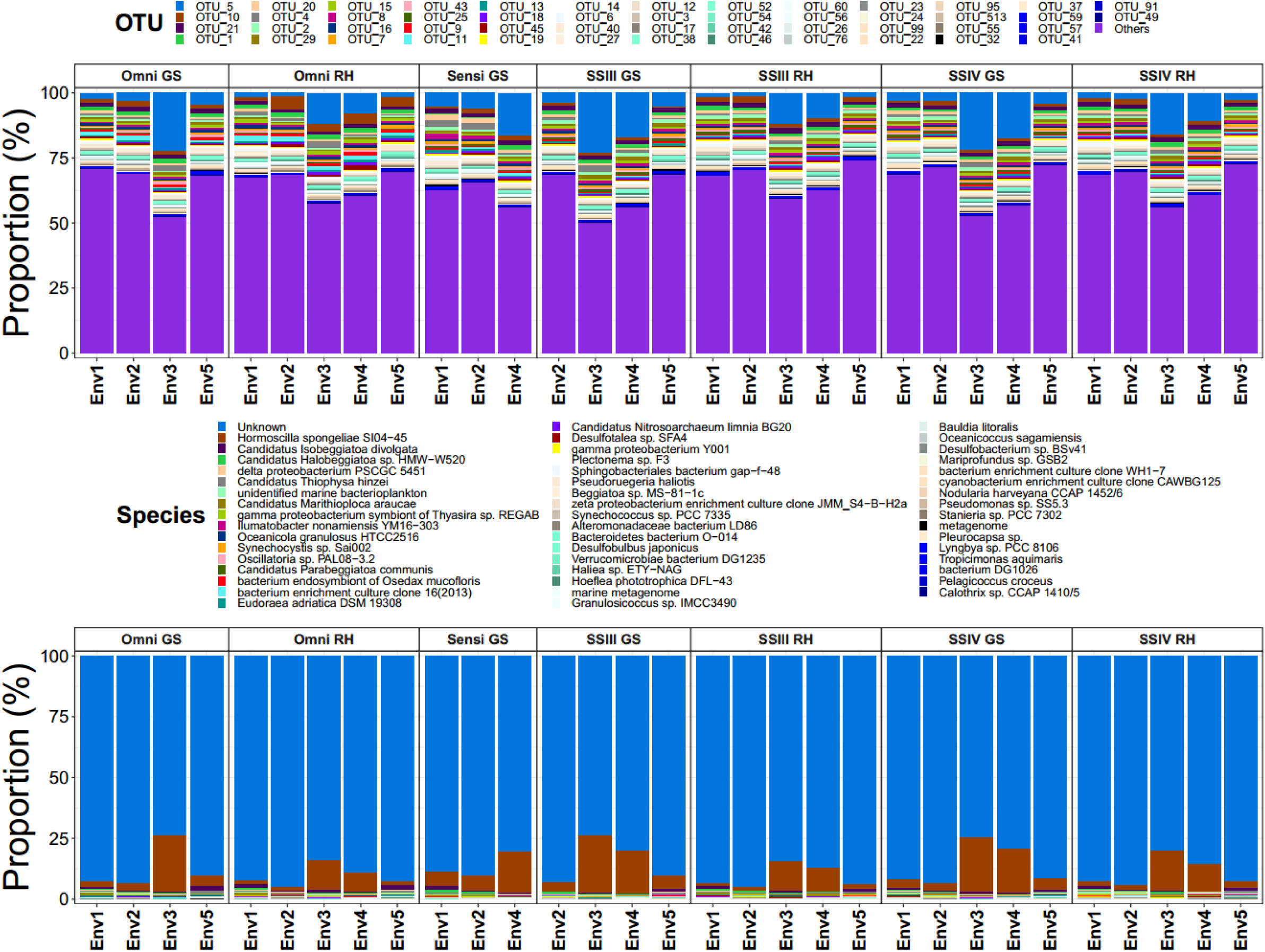
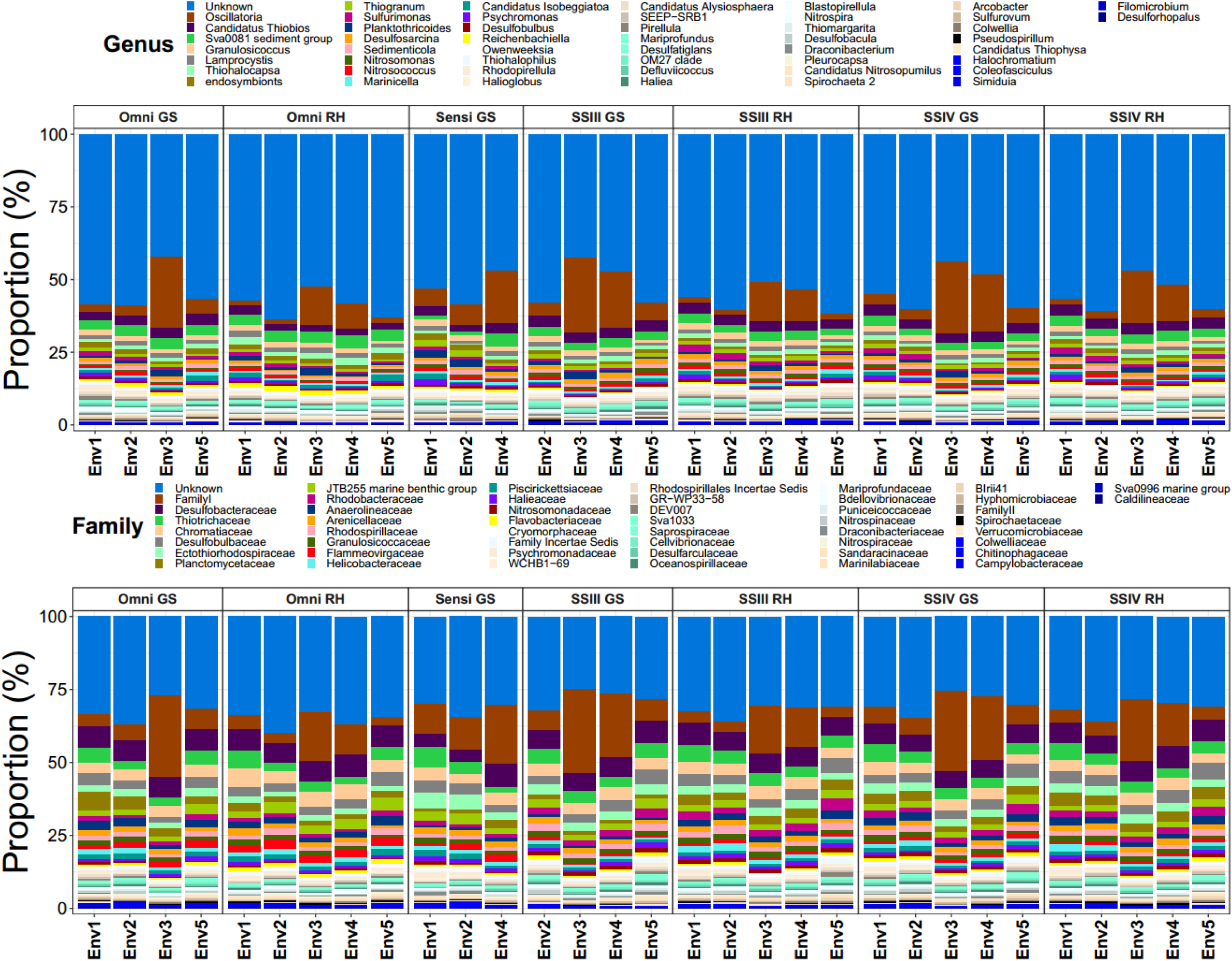
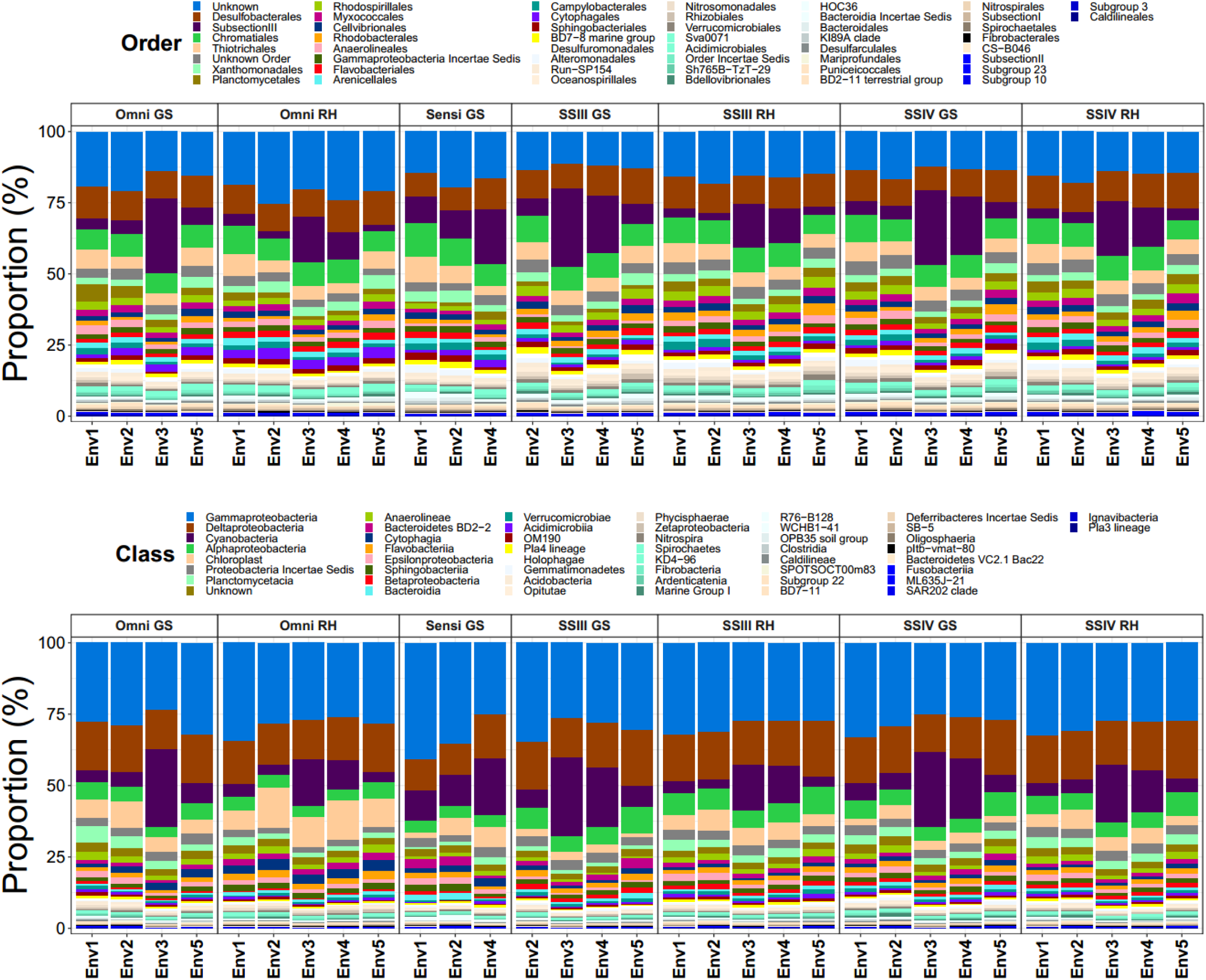
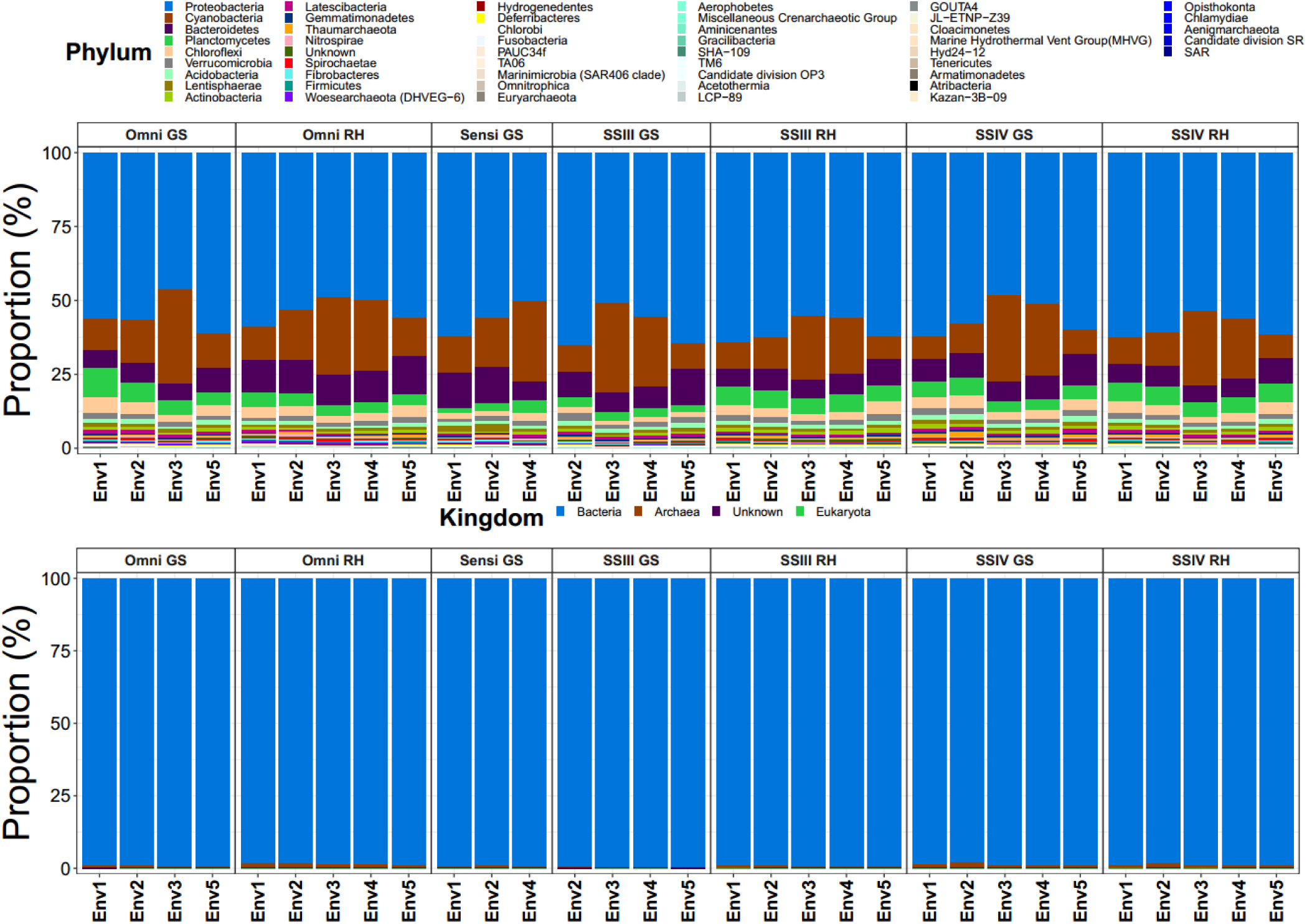
Effect of RT system on 16SrRNA transcript community composition. Bar charts represent changes in community composition of the 50 most abundant members at different taxonomic levels.

**Figure S.4.**
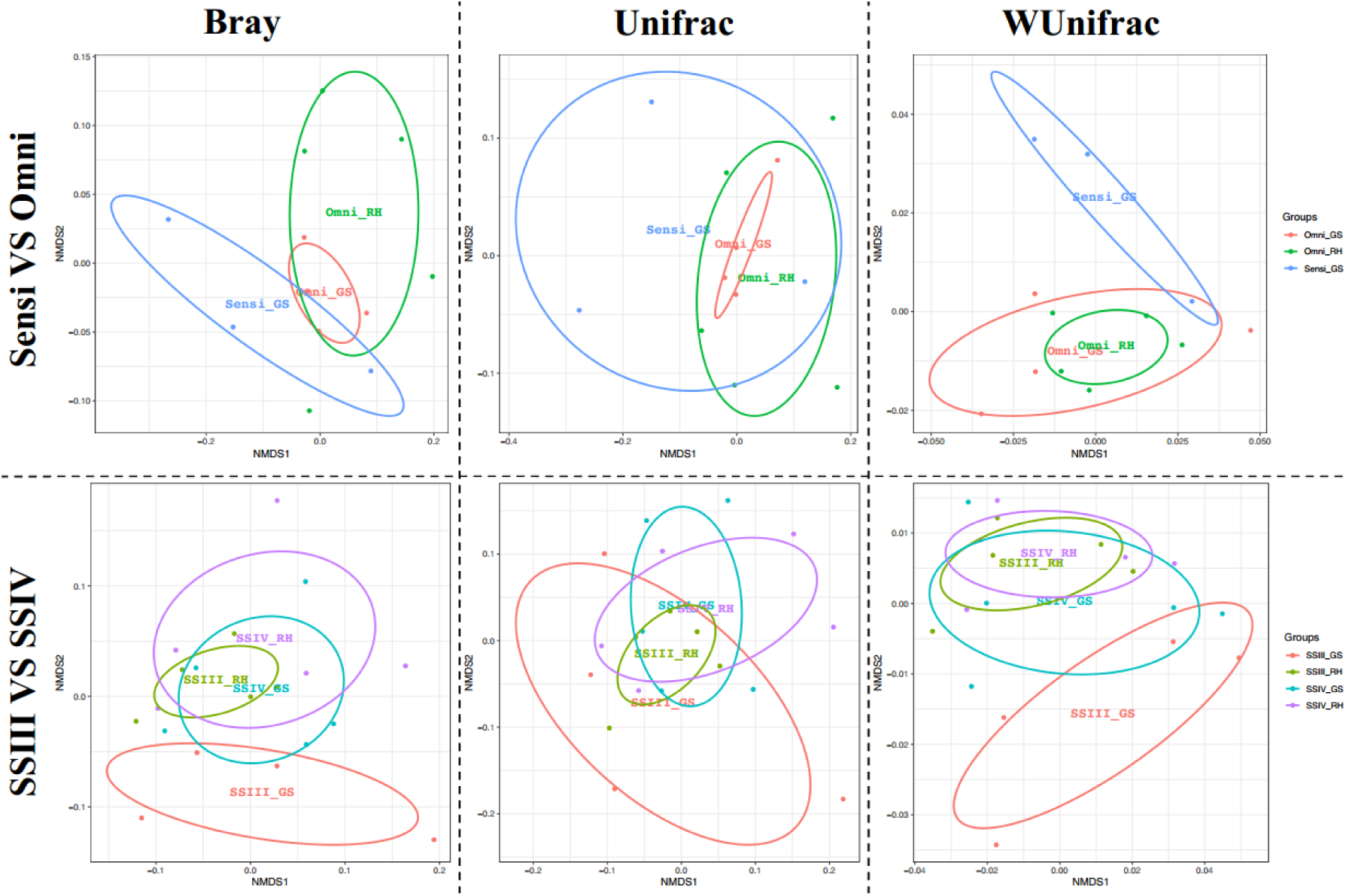
Effect of enzyme and priming on *16S rRNA* transcript community composition. NMDS clustering of *16S rRNA* cDNA community composition of the same sample derived from different enzyme and primer strategies: Sensiscript and Omniscript (top) and Superscript III and Superscript IV (Bottom). Distances were calculated using Bray-Curtis (left), Unifrac (middle) and WUnifrac (right) distances. Corresponding groups are indicated in the legend.

**Figure S.5.**
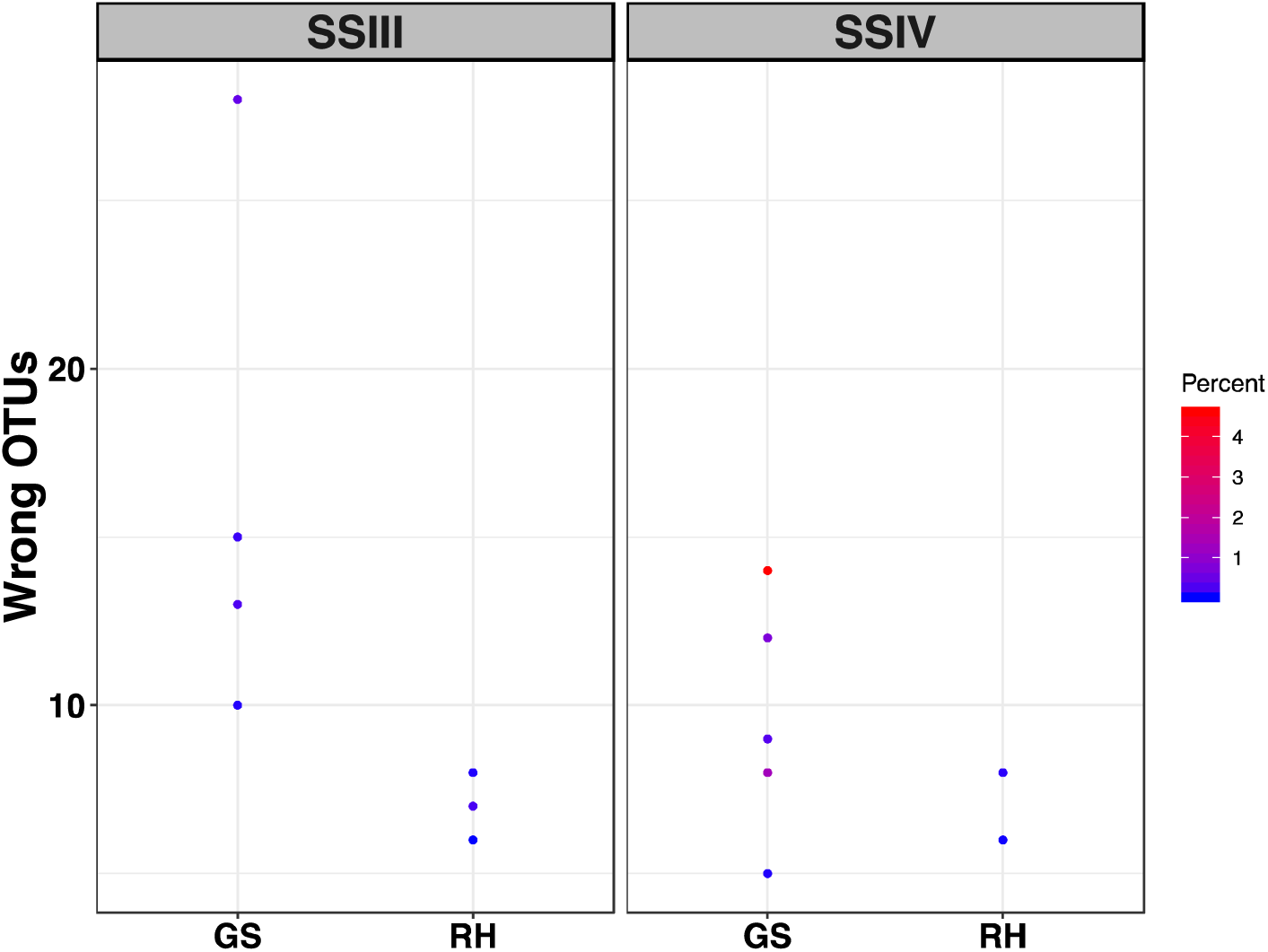
Wrong OTUs detected in the *amoA* libraries. The average number of OTUs with no assignment or wrong assignment when ran on BLASTx was plotted for each enzyme (SSIII = Superscript III; SSIV = Superscript IV) and each priming strategy (GS = Gene Specific; RH = Random Hexamer). The colour scale indicates the percentage of reads that these wrong OTUs represent compared to the total number of reads of each individual library.

**Figure S.6.**
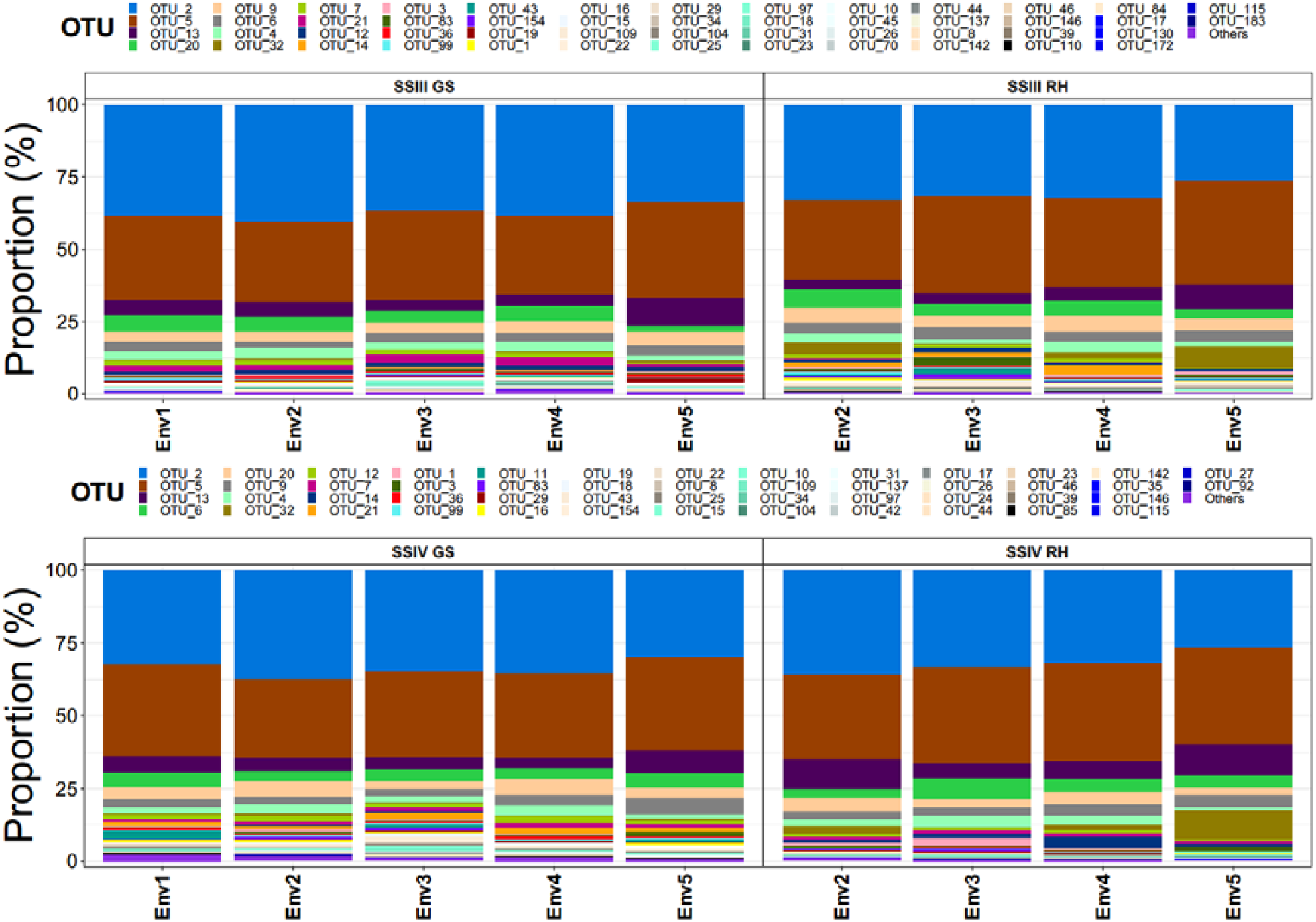
Effect of priming on amoA-OTUs transcript community composition. Bar charts represent changes in community composition of the 50 most abundant OTUs depending on the priming used for SSIII (top) and SSIV (Bottom).

**Figure S.7.**
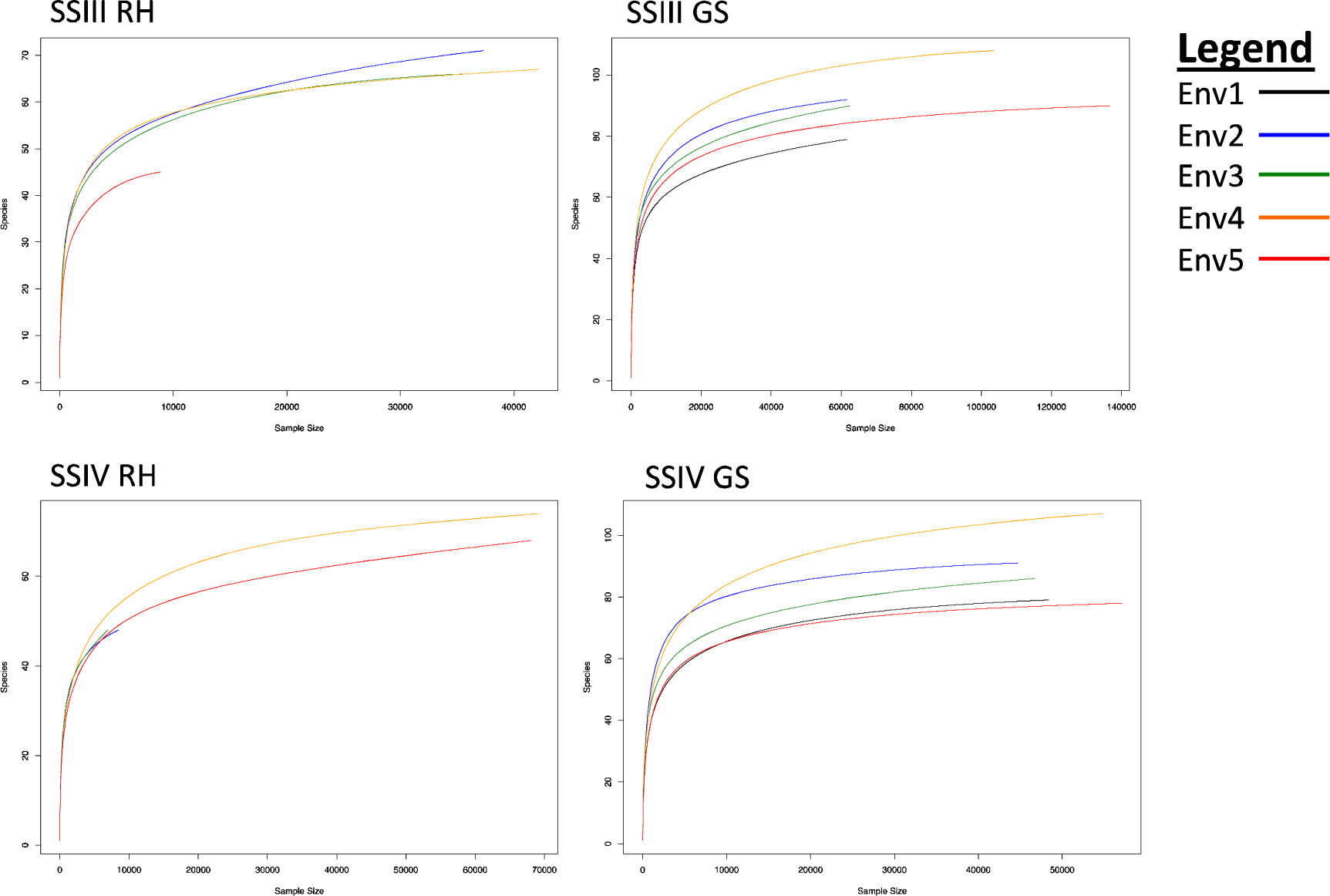
rarefaction curves obtained from the sequencing of *amoA* transcripts from environmental samples using different RT protocols. The RT protocol used is indicated on the plots (SSIII= Superscript III; SSIV= Superscript IV/ RH= random hexamer; GS= gene specific). The rarefaction curves were drawn for each replicate as indicated on the legend.

## Supplementary Tables

**Table S.1.** Description of the sfGFP standard curve regressions.

| Enzyme | Priming | Slope | Efficiency (%) | Y-Intercept | R squared | LD |
| --- | --- | --- | --- | --- | --- | --- |
| SSIV | GS | -3.34 | 99.3 | 43.55 | 0.993 | $10^3$ |
| | RH | -3.57 | 90.6 | 45.44 | 0.993 | $10^3$ |
| SSIII | GS | -3.39 | 97.2 | 43.62 | 0.996 | $10^3$ |
| | RH | -3.49 | 93.4 | 44.9 | 0.996 | $10^3$ |
| Sensi | GS | -3.61 | 89.2 | 43.87 | 0.999 | $10^3$ |
| | RH | -3.77 | 84.2 | 49.51 | 0.998 | $10^4$ |
| Omni | GS | -3.63 | 88.6 | 43.68 | 0.998 | $10^3$ |
| | RH | -3.67 | 87.3 | 46.23 | 0.997 | $10^3$ |

The regressions coefficients were calculated based on the linear model ( $y = ax + b$ ; $a$ =Slope and $b$ =Intercept). The fit of the regression is represented by the R squared values. Efficiency of the Q-PCR was calculated as Efficiency (%) = $100 \times (10^{-1/\text{slope}} - 1)$ . LD = Limit of Detection.

**Table S.2.** Comparison of models for standard curve.

| Model | Random Effect | df | Fit Criteria |  |  |  | Comparison | ChiSq | Df | Pr(>ChiSq) |
| --- | --- | --- | --- | --- | --- | --- | --- | --- | --- | --- |
|  |  |  | AIC | BIC | logLik | Deviance |  |  |  |  |
| Model1 | Intercept | 4 | 227 | 238 | -109 | 219 | 1 VS 3 | 7.38 | 2 | 0.025 |
| Model2 | Slope | 4 | 301 | 312 | -146 | 293 | 2 VS 3 | 80.86 | 2 | 2.2-16 |
| Model3 | Intercept and Slope | 6 | 224 | 240 | -106 | 212 |  |  |  |  |

For each model, the fit was assessed using four criteria namely; AIC (Akaike Information Criteria), BIC (Bayesian Information Criteria), LogLik (log Likelihood) and deviance. Models 1 and 2 were compared to model 3 and the statistic of the test is reported in the last column (Pr(>ChiSq))

**Table S.3.** **Composition of the mock communities**

| Sequence ID | Even Mock (EM) |  | Uneven Mock (UM) |  |
| --- | --- | --- | --- | --- |
|  | µl into pool | % of the community | µl into pool | % of the community |
| 1 |  |  | 0.523 | 4.36 |
| 2 |  |  | 0.407 | 3.39 |
| 3 |  |  | 1.020 | 8.57 |
| 4 |  |  | 0.880 | 7.33 |
| 5 |  |  | 0.662 | 5.52 |
| 6 | 1 | 100/12≈ 8.33 | 0.815 | 6.79 |
| 7 |  |  | 1.452 | 12.1 |
| 8 |  |  | 2.170 | 18.08 |
| 9 |  |  | 1.490 | 12.41 |
| 10 |  |  | 0.690 | 5.73 |
| 11 |  |  | 0.955 | 7.96 |
| 12 |  |  | 0.931 | 7.76 |

**Table S.4.**
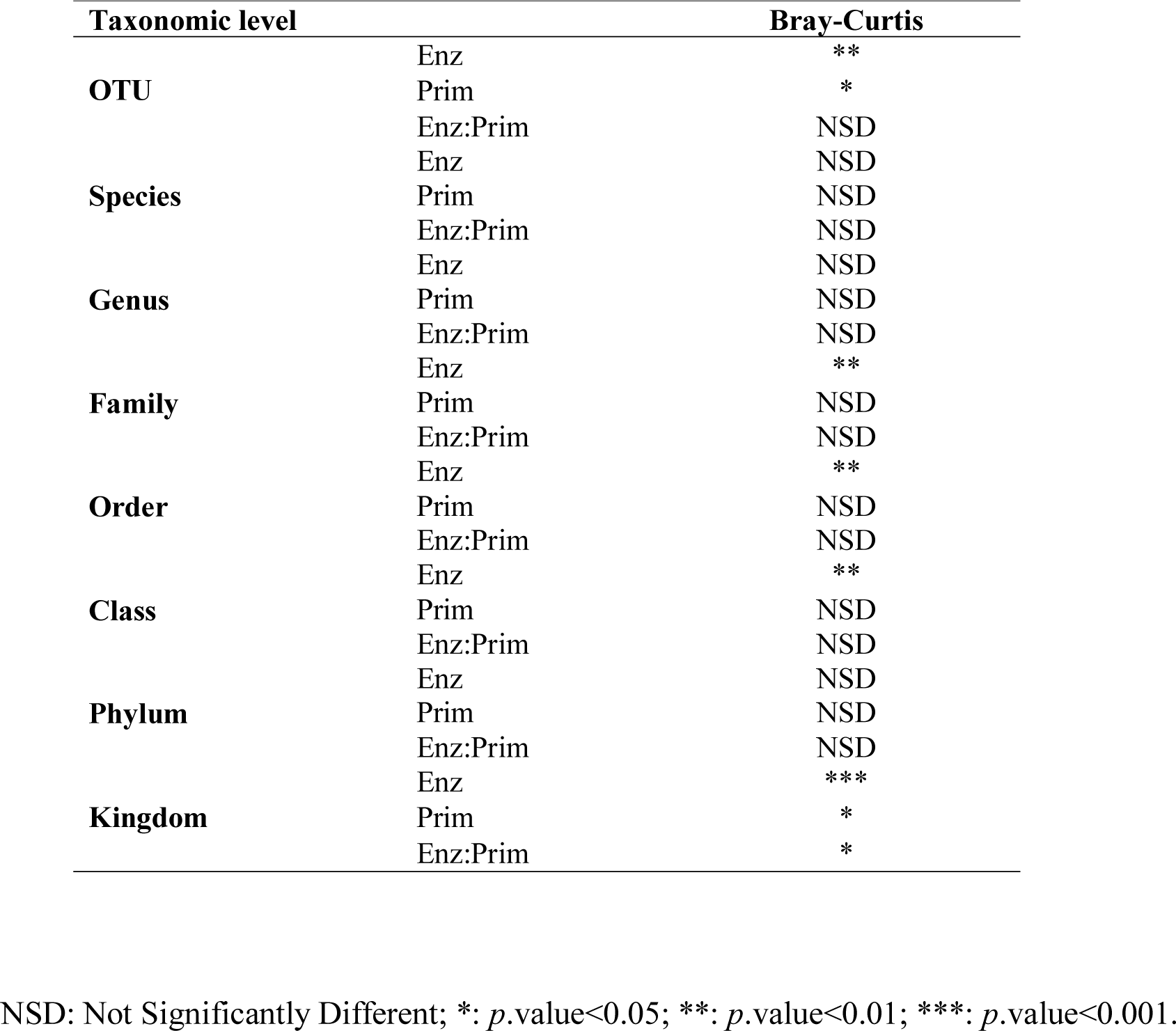
Effect of the RT system (Enzyme and Priming) on the community composition of the 16SrRNA libraries at different taxonomic levels.

| **Taxonomic level** |  | **Bray-Curtis** |
| --- | --- | --- |
| **OTU** | Enz | ** |
|  | Prim | * |
|  | Enz:Prim | NSD |
| **Species** | Enz | NSD |
|  | Prim | NSD |
|  | Enz:Prim | NSD |
| **Genus** | Enz | NSD |
|  | Prim | NSD |
|  | Enz:Prim | NSD |
| **Family** | Enz | ** |
|  | Prim | NSD |
|  | Enz:Prim | NSD |
| **Order** | Enz | ** |
|  | Prim | NSD |
|  | Enz:Prim | NSD |
| **Class** | Enz | ** |
|  | Prim | NSD |
|  | Enz:Prim | NSD |
| **Phylum** | Enz | NSD |
|  | Prim | NSD |
|  | Enz:Prim | NSD |
| **Kingdom** | Enz | *** |
|  | Prim | * |
|  | Enz:Prim | * |

NSD: Not Significantly Different; \*: $p.value < 0.05$ ; \*\*: $p.value < 0.01$ ; \*\*\*: $p.value < 0.001$ .

**Table S.5.**
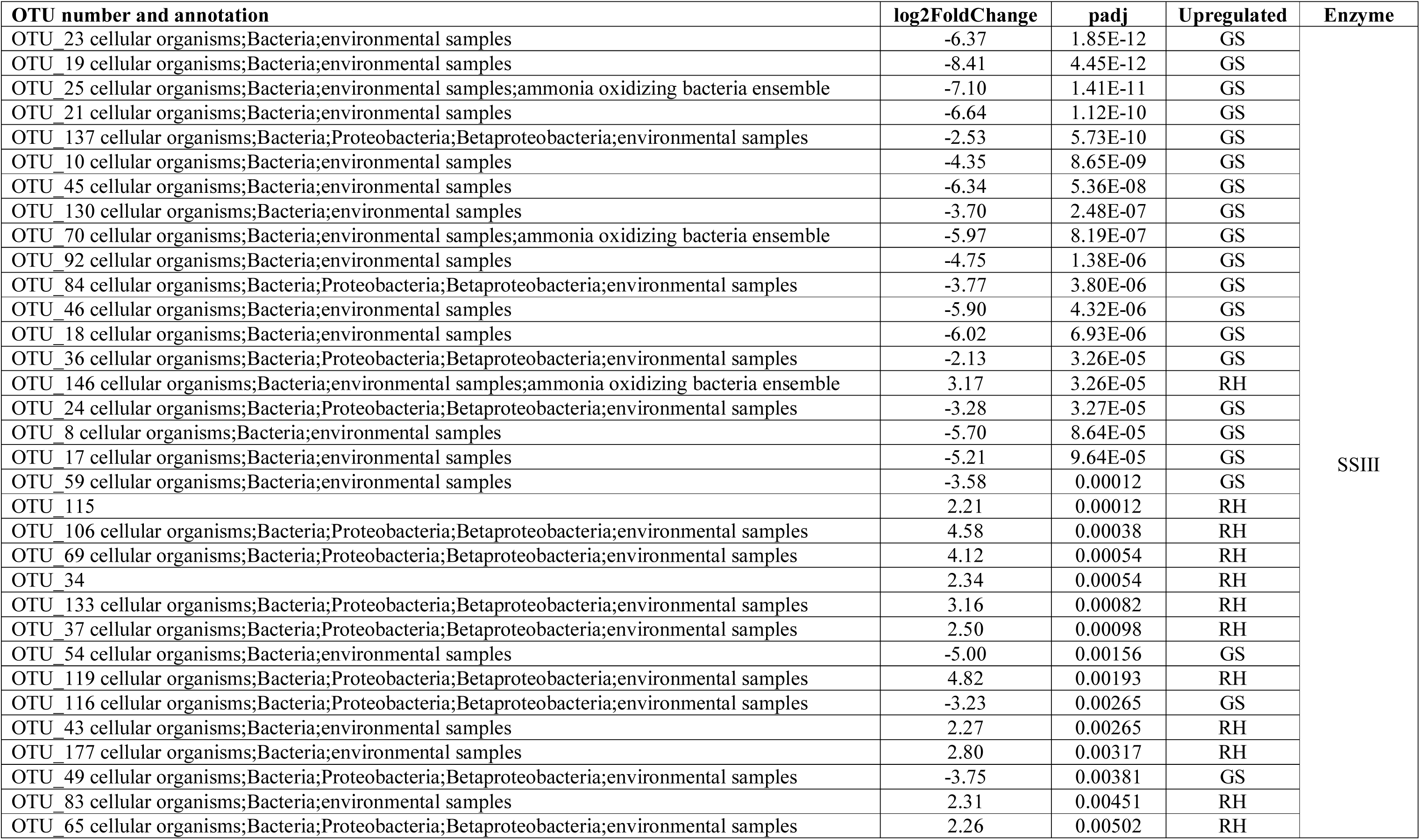
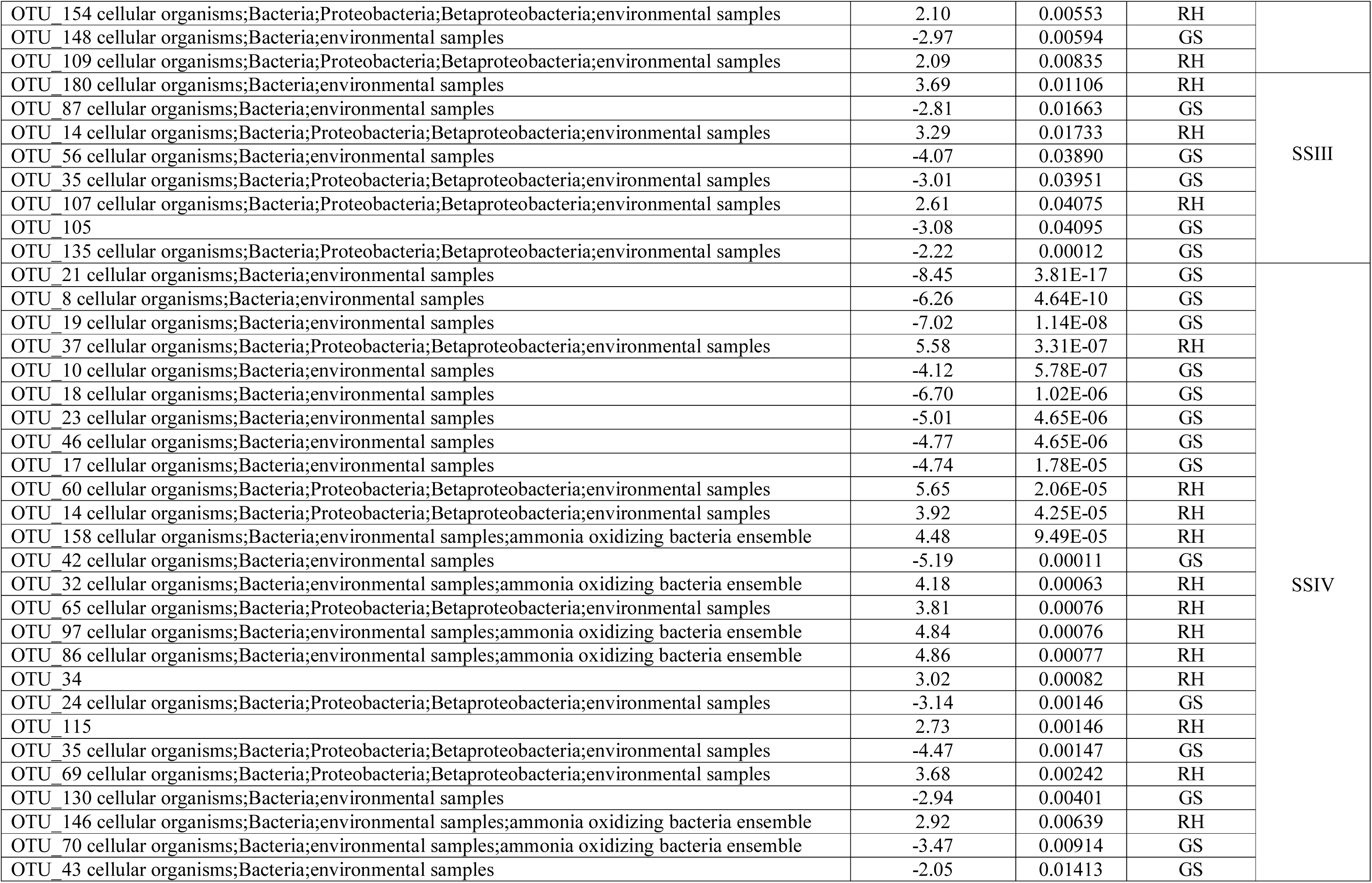
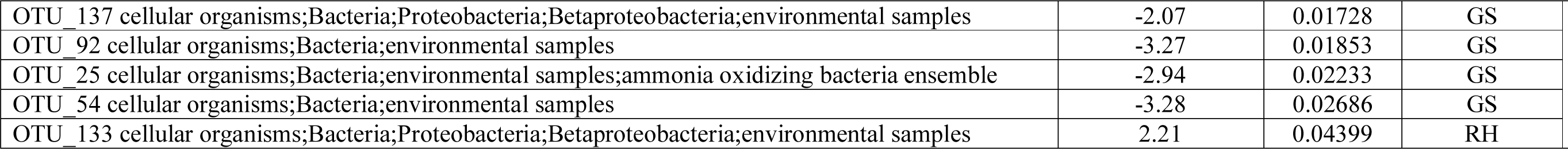
Summary of the amoA OTU for which the relative proportion is significantly different between the GS and RH databases for SSIII and SSIV

**Table S.6.**
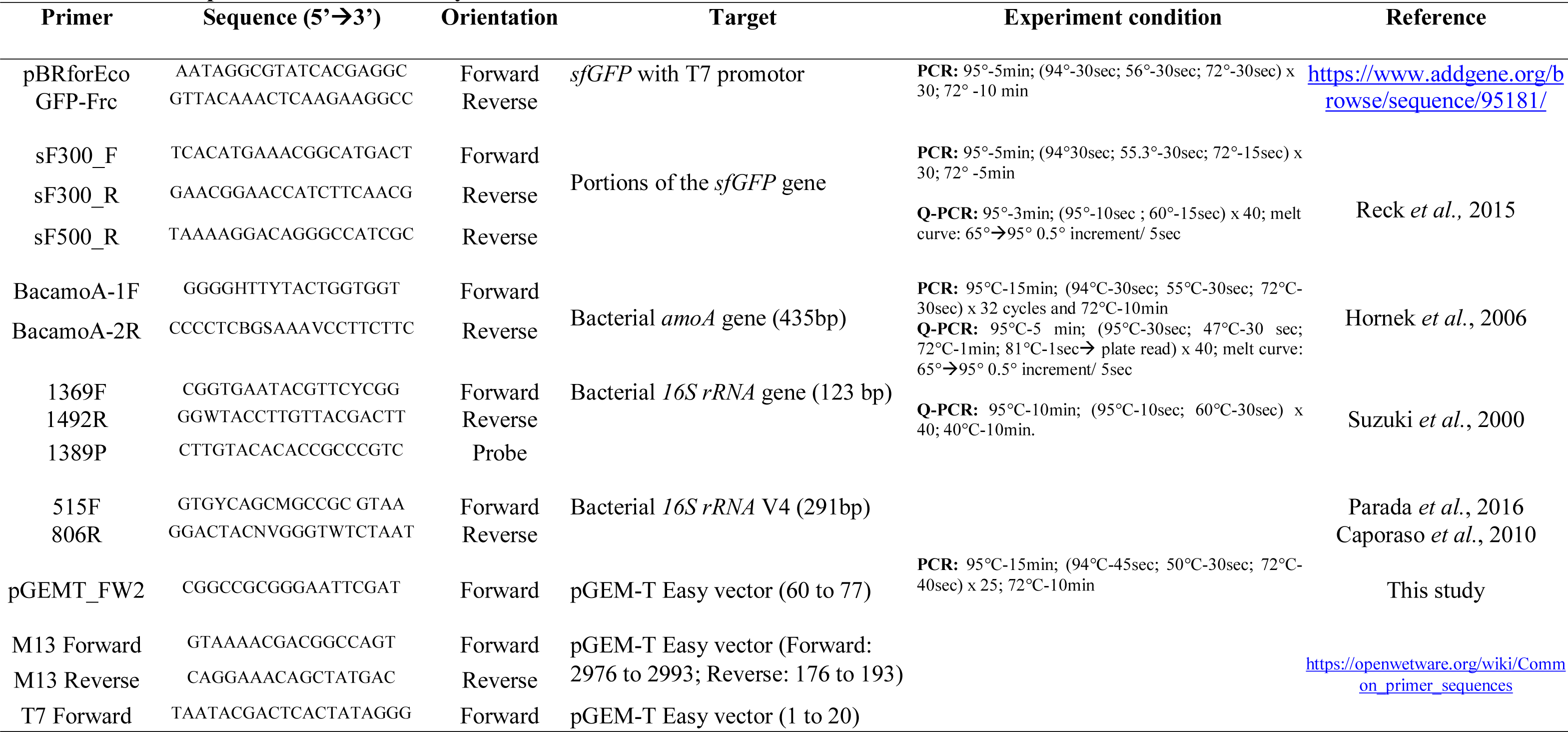
List of primers used in this study.

**Table S.7.** Reverse Transcriptase kits and reaction conditions used in this study.

| Superscript III and IV<br>(named SSIII and SSIV respectively thereafter) |  | Sensiscript and Omniscript<br>(named Omni and Sensi respectively thereafter) |
| --- | --- | --- |
| <u>Cocktail 1:</u><br>- 1µl primer (10µM GS; 50µM RH)<br>- 1µl dNTPs (10µM each)<br>- 5µl template RNA<br>- 6µl water |  | Template RNA:<br>65°C-5min; ice-1min |
| 65°-5min; ice-1min |  | <u>Cocktail:</u><br>- 2µl buffer 10x<br>- 1µl primer (10µM GS; 50µM RH)<br>- 5µl template RNA<br>- 1µM dNTPs (10µM each)<br>- 1µl enzyme (80U/µl)<br>- 1µl RNase inhibitor (40U/µl)<br>- 9µl water |
| <u>Cocktail 2:</u><br>- 4µl Buffer 5X<br>- 1µl DTT (0.1mM)<br>- 1µl enzyme (200U/µl)<br>- 1µl RNase inhibitor (40U/µl) |  |  |
| RH | GS | RH and GS |
| - 25°C (SSIII)/ 23°C(SSIV) for 10min | - 55°C for 50min (SSIII)/ 55°C for 10min (SSIV) |  |
| - 50°C for 50min (SSIII)/ 55°C for 10min (SSIV) | - 72°C for 15min (SSIII)/ 80°C for 10min (SSIV) | 37°C for 60min |
| - 72°C for 15min (SSIII)/ 80°C for 10min (SSIV) |  |  |

The black arrow indicates the procedure through time. RH = Random Hexamer; GS = Gene Specific.

**Table S.8.** Description of the *amoA* and *16SrRNA* standard curve regressions.

| Target | Enzyme | Priming | Slope | Efficiency (%) | Y-Intercept | R squared |
| --- | --- | --- | --- | --- | --- | --- |
| *amoA* | SSIV | GS | -3.30 | 100.80 | 43.68 | 0.997 |
|  |  | RH | -3.37 | 98.19 | 45.79 | 0.988 |
|  | SSIII | GS | -3.41 | 96.45 | 42.95 | 0.999 |
|  |  | RH | -3.33 | 99.78 | 44.46 | 0.996 |
|  | Omni | GS | -3.63 | 88.68 | 45.45 | 0.999 |
|  |  | RH | -3.44 | 95.39 | 47.03 | 0.982 |
|  | Sensi | GS | -3.67 | 87.26 | 46.16 | 0.994 |
|  |  | RH | -3.85 | 81.95 | 50.55 | 0.980 |
| *16S rRNA* | SSIV | GS | -3.418 | 96.14 | 43.26 | 0.995 |
|  |  | RH | -3.423 | 95.95 | 44.19 | 0.969 |
|  | SSIII | GS | -3.16 | 107.38 | 42.00 | 0.997 |
|  |  | RH | -3.49 | 93.47 | 44.45 | 0.991 |
|  | Omni | GS | -3.10 | 110.12 | 41.93 | 0.994 |
|  |  | RH | -3.03 | 113.81 | 47.97 | 0.9535 |
|  | Sensi | GS | -2.921 | 119.96 | 41.215 | 0.9912 |
|  |  | RH | -2.78 | 128.94 | 43.7 | 0.9978 |

## Notes

http://www.ebi.ac.uk/ena/data/view/PRJEB32314

